# Entry to and exit from diapause arrest in *Caenorhabditis elegans* are both regulated by a steroid hormone pathway

**DOI:** 10.1101/2021.09.04.458989

**Authors:** Mark G Zhang, Paul W Sternberg

## Abstract

Diapause arrest in animals such as *Caenorhabditis elegans* is tightly regulated so that animals make appropriate developmental decisions amidst environmental challenges. Fully understanding diapause requires mechanistic insight of both entry and exit from the arrested state. While a steroid hormone pathway regulates the entry decision into *Caenorhabditis elegans* dauer diapause, its role in the exit decision is less clear. A complication to understanding steroid hormonal regulation of dauer has been the peculiar fact that steroid hormone mutants such as *daf-9* form partial dauers under normal growth conditions. Here, we corroborate previous findings that *daf-9* mutants remain capable of forming full dauers under unfavorable growth conditions, and we establish that the *daf-9* partial dauer state is likely a partially exited dauer that has initiated but cannot complete the dauer exit decision. We show that the steroid hormone pathway is both necessary for and promotes complete dauer exit, and that the spatiotemporal dynamics of steroid hormone regulation during dauer exit resembles that of dauer entry. Overall, dauer entry and dauer exit are distinct developmental decisions that are both controlled by steroid hormone signaling.

**Summary Statement:** In animals such as *Caenorhabditis elegans*, a steroid hormone pathway controls both the entry and exit decisions into and out of the developmentally arrested dauer state in response to environmental signaling.

## Introduction

Animals must be able to adapt to changing environments to survive against uncertain and stress-inducing circumstances. One such adaptive mechanism is diapause, a state of developmental arrest typically characterized by metabolic depression and stress resistance (Hand et al., 2016). Diapause is a dynamic process that involves successive developmental decisions dictating entry, maintenance, and exit from the dormant state (Koštál, 2006; Ragland et al., 2010). Diapause is well conserved across the animal kingdom including nematodes, insects, crustaceans, fish, and mammals (Hand et al., 2016; Hu, 2007; Podrabsky and Hand, 2015; Ragland et al., 2010).

Upon encountering adverse conditions during larval growth, *Caenorhabditis elegans* exit the cycle of reproductive development and instead enter the alternative diapause state, termed dauer, which grants them increased durability and longevity to protect against environmental insults and allows them to disperse in search of a more favorable environment (Cassada and Russell, 1975; Hu, 2007). The dauer entry decision-making process comprises two distinct subdecisions. First-stage (L1) larvae decide between developing into L2 or pre-dauer L2d larvae, depending on whether conditions are favorable or unfavorable, respectively (the “L1 to L2/L2d subdecision”). If conditions sufficiently improve, then L2d larvae choose reproductive development by becoming L3 larvae, but if not they become dauer larvae (the “L2d to L3/Dauer subdecision”; Golden and Riddle, 1984). While in the dauer state, animals continuously assess their environment and, when conditions improve by way of an increased food to pheromone ratio, exit the dauer state to return to the reproductive cycle as L4 larvae (Golden and Riddle, 1982). The *C. elegans* dauer process thus involves two consecutive entry subdecisions into the dauer state as well as the exit decision to return reproductive development. A complete understanding of this developmental decision-making process requires a synthesis of information involving both the dauer entry subdecisions and the dauer exit decision. The vast majority of dauer studies in *C. elegans* have focused almost exclusively on dauer entry (Androwski et al., 2017; Fielenbach and Antebi, 2008; Hu, 2007), leaving much to be explored for dauer exit.

Previous studies have found multiple pathways that govern the dauer entry decision, including cGMP signaling, insulin growth factor signaling, TGF-β signaling, and steroid hormone signaling (Fielenbach and Antebi, 2008; Hu, 2007). Early genetic screens for dauer exit defects yielded identical genes to those involved in dauer entry, suggesting that the same genetic pathways underlie both the dauer entry and exit decisions (Riddle et al., 1981). But even though the same principal pathways might govern both decisions, a wealth of evidence suggests important distinctions exist between the dauer entry and exit decision, including the involvement of different sensory neurons (Bargmann and Horvitz, 1991), the different timeframes of the decision-making process (Golden and Riddle, 1984; Schaedel et al., 2012), and the differing importance of various signaling molecules such as the insulin peptides and the Notch ligands (Cornils et al., 2011; Ouellet et al., 2008). These differences highlight the importance of studying the dauer exit decision in tandem with the dauer entry decisions in order to construct a comprehensive regulatory model of the dauer process.

Steroid hormones have been characterized as one of the most downstream components in the dauer entry process and are thought to serve as a convergence point for both the insulin and the TGF-β pathway (Fielenbach and Antebi, 2008) in controlling dauer development. The steroid hormone pathway centers on DAF-12, a nuclear hormone receptor that shows the most homology to the vertebrate farnesoid-X receptor (FXR) (Antebi, 2015; Antebi et al., 1998). The major endogenous ligands for DAF-12/FXR are steroid hormones collectively referred to as dafachronic acids (DA), which include Δ7-DA (dafa#2) and Δ4-DA (dafa#4) among others (Aguilaniu et al., 2016; Mahanti et al., 2014; Motola et al., 2006). DAF-12/FXR regulation of its transcriptional targets depends on environmental growth conditions, which in turn dictate the presence of DAF-12/FXR ligands. Under favorable conditions, DAF-9 catalyzes the formation of DAs such as Δ-7 DA which bind to DAF-12/FXR and specify reproductive adulthood. *daf-12* alleles with mutations in the putative ligand binding domain are therefore mostly <u>da</u>uer <u>f</u>orming <u>c</u>onstitutive (Daf-c) (Antebi et al., 1998). Under unfavorable conditions, unliganded DAF-12/FXR interacts with the corepressor DIN-1/CoR to specify dauer entry (Fielenbach and Antebi, 2008). *daf-12* null alleles are thus <u>da</u>uer <u>f</u>ormation <u>d</u>efective (Daf-d).

Biosynthesis of all known DAs require the cytochrome P450 enzyme DAF-9, and therefore *daf-9* null mutants are completely Daf-c (Aguilaniu et al., 2016; Gerisch et al., 2001; Jia et al., 2002). *daf-9* is constitutively expressed in a pair of neuroendocrine cells termed the XXX cells but show variable upregulation in the hypoderm depending on environmental conditions and developmental state (Gerisch et al., 2001; Schaedel et al., 2012). During the L2d to L3/dauer subdecision, sufficient exposure to favorable conditions causes upregulation of *daf-9* throughout the hypoderm that amplifies steroid hormone production to instigate reproductive development (Gerisch et al., 2001; Schaedel et al., 2012). Hypodermal expression of *daf-9* suffices to bypass dauer in *daf-9* animals, suggesting that the hypoderm is an important site of *daf-9* expression (Gerisch and Antebi, 2004). Here, we sought to characterize how steroid hormonal regulation of dauer exit compares to its role in the two dauer entry subdecisions.

A complete analysis concerning how steroid hormone signaling regulates dauer arrest must also account for the well-documented observation that Daf-c mutants impaired for steroid hormone biosynthesis and/or signaling such as *daf-9, daf-36* (encoding a Rieske-like oxygenase that catalyzes the first step of steroid hormone biosynthesis, *ncr-1; ncr-2* (encoding two putative cholesterol transporters), and Daf-c alleles of *daf-12* do not form full dauers under normal growth conditions as do other Daf-c strains such as *daf-2(e1370)* or *daf-7(e1372)* (Antebi et al., 1998; Gerisch et al., 2001; Li et al., 2004; Rottiers et al., 2006). Full dauers are characterized by radial and pharyngeal constriction, immobility, pumping quiescence, and a darkened intestine owing to increased fat storage (Cassada and Russell, 1975; Riddle and Albert, 1997). Steroid hormone mutants such as *daf-9* instead form “partial” or “dauer-like” larvae that resemble dauers but exhibit non-dauer traits such as sporadic pumping, increased mobility, a slightly enlarged pharynx, and a lighter body (Albert and Riddle, 1988; Gerisch et al., 2001). Partial dauers are not exclusive to steroid hormone mutants, as they are also observed in double mutants involving *daf-16* (Ailion and Thomas, 2000; Vowels and Thomas, 1992), which encodes a homolog of the forkhead transcription factor FOXO and controls the insulin pathway (Ogg et al., 1997).

Whether steroid hormone mutants form partial or full dauers appear to depend on growth conditions. Anecdotal evidence indicates that *daf-9(dh6)* and *daf-12(rh273)* become full dauers under unfavorable growth conditions (Antebi et al., 1998; Gerisch et al., 2001). Why this distinction occurs remains unclear, but it has been speculated that the partial dauer may have first been a full dauer that attempted dauer exit (owing to favorable growth conditions) but could not complete it (Antebi et al., 1998). A comprehensive model of how steroid hormones govern dauer entry and exit should be able to explain why steroid hormone mutants form partial dauers under favorable growth conditions unlike other Daf-c strains (**Fig. 1**).

**Figure 1.**
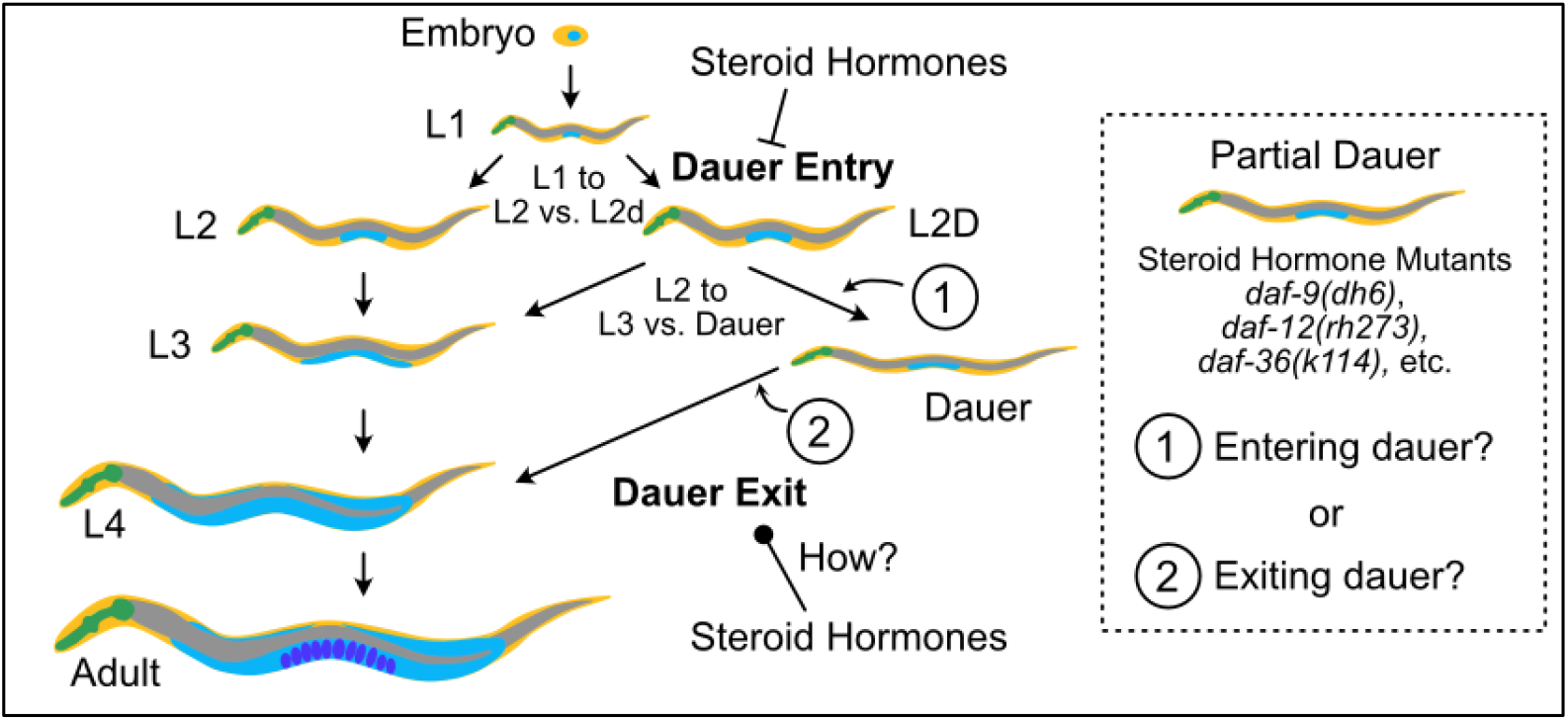
Regulation of the *Caenorhabditis elegans* dauer developmental pathway by steroid hormones. The *C. elegans* dauer pathway includes multiple developmental decisions. The dauer entry decision comprises two subdecisions made at L1 and then L2d, whereas dauer exit is a singular, continuous decision. Leftward or rightward arrows indicate the decision that is made under favorable or unfavorable conditions, respectively. Steroid hormones inhibit dauer entry and promote reproductive development. How steroid hormones regulate dauer exit is less understood. A model of how steroid hormones regulate the dauer pathway requires understanding why steroid hormone mutants such as *daf-9(dh6), daf-12(rh273), daf-36(k114)*, etc. form partial dauers rather than full dauers like other Daf-c mutants. Partial dauers could be (1) animals that have yet to enter a full dauer state or (2) partially exited dauers.

In this study, we provide evidence that *daf-9* partial dauers are indeed animals that first became full dauers that then initiated (but failed to complete) dauer exit. Furthermore, we find that the *daf-9* partial dauer state is dependent on *daf-2* and *daf-7*, suggesting that the partial dauer state arises due to activation of the insulin or the TGF-β pathway. We characterize the regulatory role of steroid hormones in dauer exit, demonstrating that the steroid hormone biosynthesis pathway is both necessary for and sufficient to induce dauer exit. We also show that the spatiotemporal regulation of *daf-9* during dauer exit closely mirrors that of the L1 to L2 vs. L2d dauer entry subdecision. Our findings reveal that *C. elegans* use steroid hormone signaling in similar ways to regulate both the dauer entry and exit developmental decisions.

## Results

### *daf-9* mutants form full dauers under unfavorable growth conditions

To assess the role of steroid hormones in the *C. elegans* dauer exit developmental decision (**Fig. 1**), we chose to focus on *daf-9* since it is the only steroid hormone gene for which null mutants such as *daf-9(dh6)* and *daf-9(e1406)* show completely penetrant Daf-c phenotypes, suggesting that its loss severely abrogates steroid hormone signaling (Antebi, 2015; Gerisch and Antebi, 2004; Gerisch et al., 2001; Rottiers et al., 2006). We first confirmed that *daf-9* null mutants could form full dauers that would be suitable for subsequent dauer exit analysis. Only anecdotal evidence suggests that the Daf-c steroid hormone mutants *daf-9(dh6)* and *daf-12(rh273)* form full dauers under unfavorable growth conditions (Antebi et al., 1998; Gerisch et al., 2001). To test whether steroid hormone mutants retain the ability to form full dauers, we grew *daf-9(dh6)* animals under unfavorable conditions, which involves high temperature (25.5°C) and the presence of dauer-inducing pheromone extract (see Materials and Methods). These unfavorable growth conditions yielded *daf-9(dh6)* dauer larvae that matched the characteristics of full dauers formed by wild-type animals. Namely, both *daf-9(dh6)* and *daf-9(e1406)* dauers showed no pumping, low motility, and a darkened, radially constricted body (**Fig. 2** and **Video 1** for *daf-9(dh6)*, **Fig. 3E-F** for *daf-9(e1406)*). *daf-9(dh6)* full dauers are also completely SDS resistant – a hallmark of the dauer state (**Fig. S1A**). In contrast, when *daf-9(dh6)* mutants were grown under favorable conditions, we observed partial dauers that pumped frequently, moved faster, and showed enlarged pharynxes compared to full dauers (**Fig. 2**). Hereafter, we distinguish between “full dauers” (which match the phenotype of wild-type dauers formed under dauer-inducing conditions) and “partial dauers” or “dauer-like larvae” (which we and others have observed when *daf-9* mutants are grown under favorable conditions).

**Figure 2.**
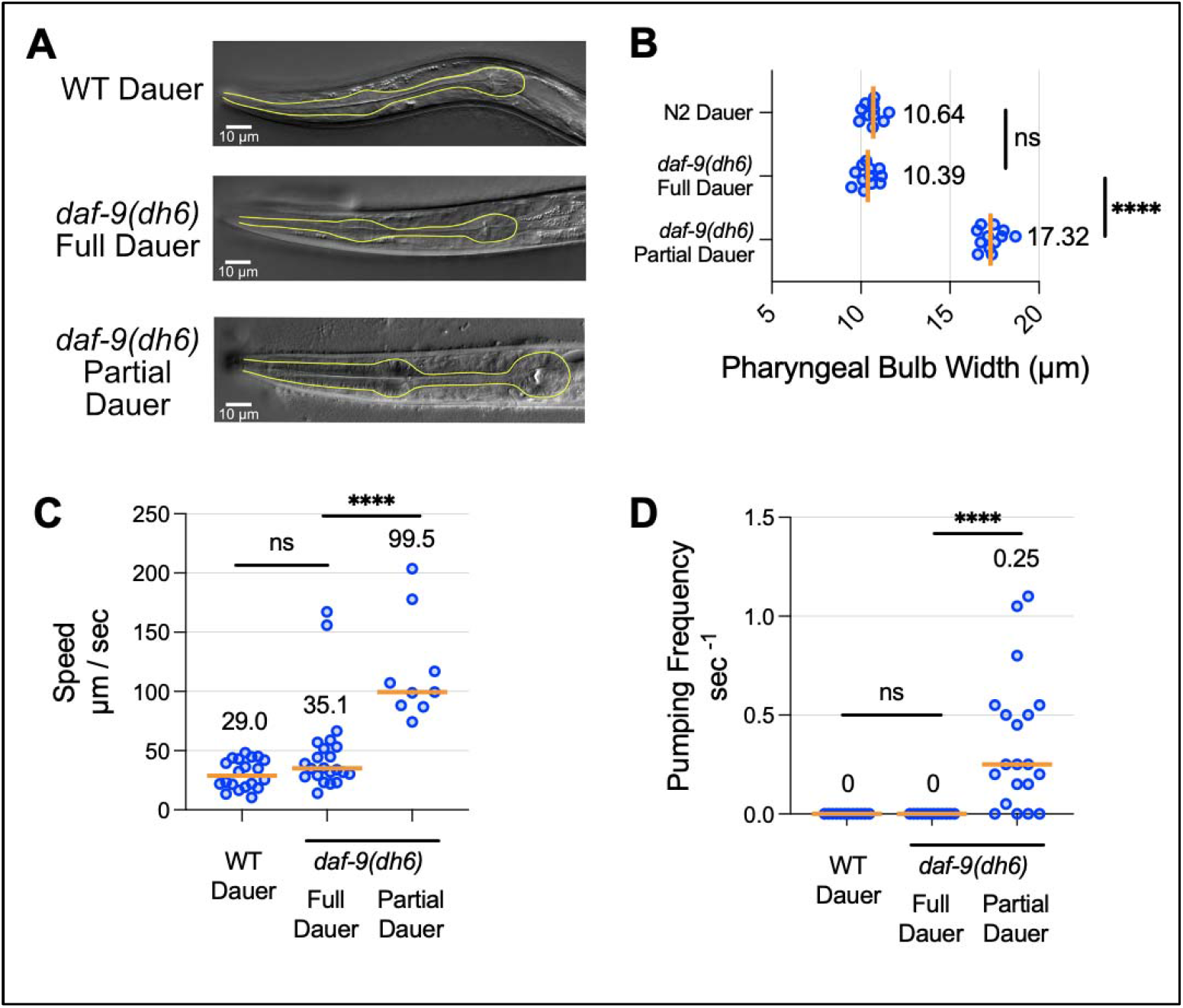
Characterization of full versus partial dauers formed by *daf-9* null mutants. *daf-9(dh6)* full dauers formed under unfavorable growth conditions phenocopy wild-type dauers and are distinct from *daf-9(dh6)* partial dauers formed under favorable growth conditions. (A) DIC images show that *daf-9(dh6)* full dauers have the same pharyngeal shrinkage as wild-type dauers, while *daf-9(dh6)* partial dauers have an enlarged pharynx. Pharynxes are outlined in yellow. (B) Width measurements of terminal pharyngeal bulbs quantifying size differences shown in (A). (C,D) *daf-9(dh6)* full dauers phenocopy wild-type dauers in terms of (C) movement speed and (D) pumping quiescence, while *daf-9(dh6)* partial dauers move more and pump more frequently. ns, not significant. ****, p<0.0001 by Mann Whitney Test. Each dot is one animal, and n ≥ 9 animals for all experimental conditions. Horizontal bars and in-graph numbers show median values.

**Figure 3:**
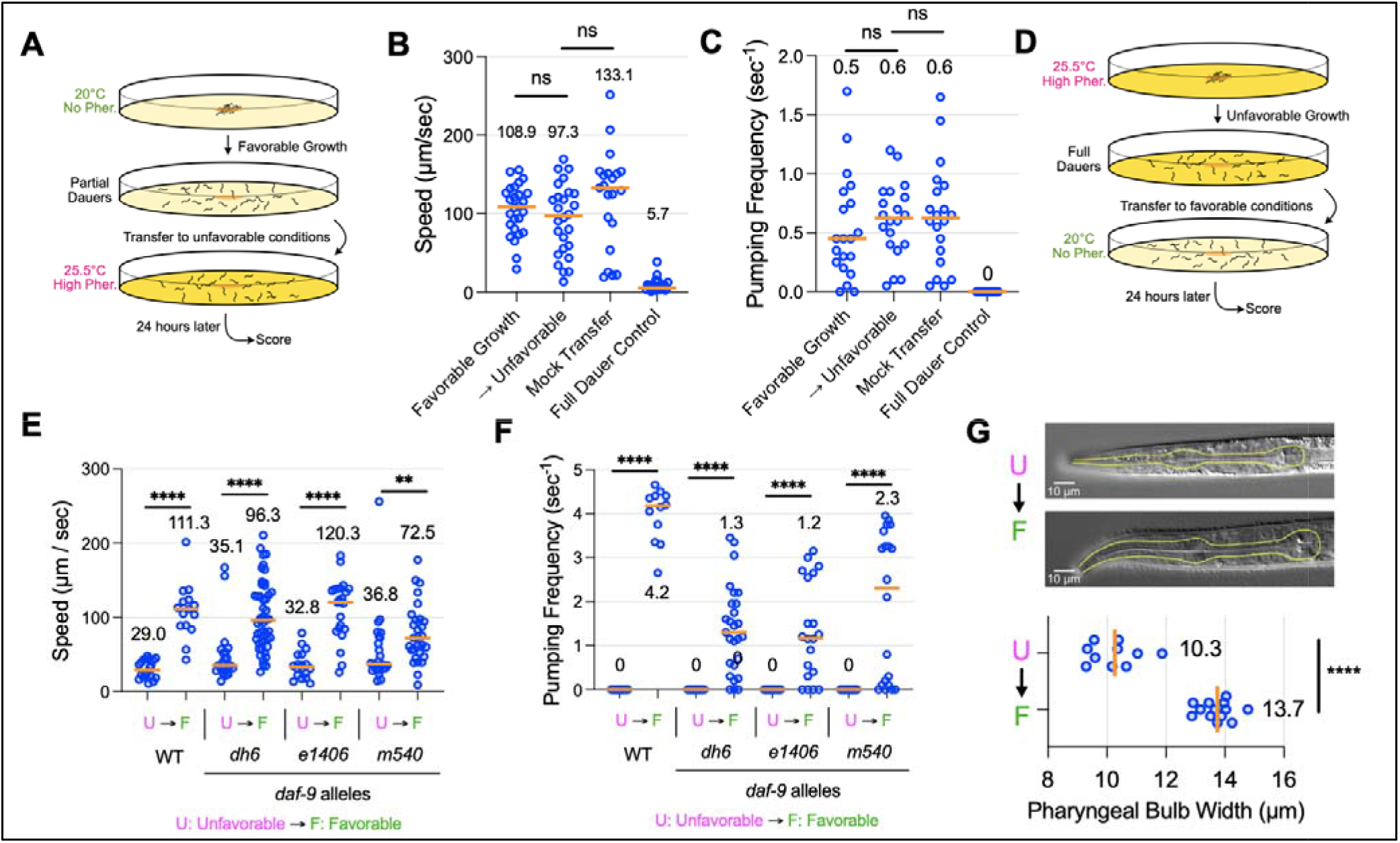
*daf-9* partial dauers may be partially exited dauers. (A) Experimental schematic to test whether *daf-9(dh6)* partial dauers become full dauers upon transfer t**o** unfavorable conditions. *daf-9(dh6)* partial dauers obtained under favorable growth conditions were transferred to unfavorable conditions or again to favorable conditions (mock transfer) and scored 24 hours later for (B) speed and (C) pumping frequency. Also shown are *daf-9(dh6)* full dauers grown under unfavorable conditions (Full Dauer Control). (D) Experimental schematic to test whether *daf-9(dh6)* full dauers become partial dauers upon transfer to favorable growth conditions. *daf-9(dh6)* full dauers were obtained under unfavorable, dauer-inducing growth conditions, transferred to favorable conditions, and then scored 24 hours later for (E) speed and (F) pumping frequency. *daf-9* alleles *e1406* (null) and *m540* (weak loss-of-function) show similar partial dauer exit phenotypes. (G) *daf-9(dh6)* partially exited dauers obtained via the method in (D) have wider pharynxes than their full dauer counterparts. ns, not significant. **, p<0.01. ****, p<0.0001 by Mann Whitney test. Each dot is one animal, and n ≥ 10 animals for all experimental conditions. Horizontal bars and in-graph numbers show median values.

### *daf-9* partial dauers are likely partially exited dauers

Having confirmed that *daf-9* mutants form partial or full dauers depending on the environmental conditions, we wanted to further probe the robust *daf-9* partial dauer phenotype, reasoning that it would provide insights into understanding how steroid hormones regulate the dauer process. Based on speculations from previous reports, we reasoned that two likely possibilities for how *daf-9* partial dauers fit into the developmental pathway are that (1) partial dauers are *en route* to becoming full dauers but require unfavorable environmental stimuli to complete the dauer entry process, or (2) partial dauers are partially exited dauers that have gone through a full dauer state and then initiate, but cannot complete, dauer exit (**Fig. 1A**). A third possibility could also be partial dauers do not fit neatly into the existing developmental pathway, and that perhaps they bypass the full dauer state completely.

To test possibility (1), in which *daf-9* partial dauers require unfavorable conditions to become full dauers, we grew *daf-9(dh6)* mutants under favorable conditions to first form partial dauers and then transferred them to unfavorable conditions to determine if they could form WT-dauers (**Fig. 3A-C**). We found that, despite a 24-hour incubation under unfavorable conditions, *daf-9(dh6)* partial dauers did not transition towards a full dauer state. These *daf-9(dh6)* animals continued to move and pump at high rates in comparison to *daf-9(dh6)* full dauers. Thus, we find it unlikely that the partial dauer state obtained under favorable growth conditions represents a transition state that is *en route* to becoming full dauer.

To assess possibility (2), in which *daf-9* partial dauers are first full dauers that then partially exit dauer, we grew *daf-9(dh6)* mutants under unfavorable conditions to first form full dauers, and then we transferred them to favorable conditions to examine if they became partial dauers. We found that 24 hours post-transfer, *daf-9(dh6)* larvae actively pumped, moved significantly more, and had an enlarged pharynx compared to before the transfer (**Fig. 3D-G,** and **Video 2**), thereby recapitulating the *daf-9* partial dauer state. These partially exited dauers slowly continued to grow radially and develop a larger pharynx even past the 24-hour mark, although they never develop into healthy reproductive adults (data not shown). We obtained similar results using *daf-9* alleles *e1406* (another putative null mutation) and *m540* (a weaker loss-of-function allele) (**Fig. 3E, F**). From both mutants, we obtained full dauers under unfavorable growth conditions that became partial dauers following transfer to favorable conditions. The major allele-specific difference was that *daf-9(m540)*, which only has a mild Daf-c phenotype, was sometimes able to form healthily exited dauers rather than partially exited dauers upon returning to favorable growth conditions (see **Fig. 3F**, *m540* pumping frequency). Together, these findings suggest that transfer to favorable conditions causes full *daf-9* dauers to initiate dauer exit and engage in concomitant behavioral and morphological changes such as increased pumping, motility, and pharyngeal expansion. Strong loss-of-function mutants such as *daf-9(dh6)* and *daf-9(e1406)* are halted in this exit process, while weaker loss-of-function mutants like *daf-9(m540)* can proceed to full reproductive adulthood.

We also examined whether this partial dauer exit phenotype could be recapitulated at the level of the nuclear hormone receptor DAF-12/FXR, which acts genetically after DAF-9 to specify the dauer decision. Although most loss-of-function alleles for *daf-12* are Daf-d, there exists numerous Daf-c alleles that bear mutations altering DAF-12’s putative ligand binding domain (Antebi et al., 1998, 2000). These Daf-c strains have also been described to form partial dauers under favorable growth conditions, although at a lower penetrance than *daf-9* putative null mutants (Antebi et al., 1998). We took such a Daf-c mutant, *daf-12(rh273)*, and found that we were able to induce full dauers under unfavorable growth conditions that could become partial dauers upon transfer to favorable conditions (**Fig. S1B, C**). Therefore, *daf-12(rh273)* mutants can phenocopy the partial dauer exit phenotype of *daf-9* putative null mutants, consistent with DAF-12/FXR mediating this phenotype.

A feature of dauer exit is its irreversibility: wild-type dauers that have been shifted to favorable conditions commit to dauer exit within an hour since shifting them back onto unfavorable conditions afterwards cannot maintain or restore the dauer state (Golden and Riddle, 1984). We asked whether the *daf-9* dauer-like state represented a committed or irreversible state of partial dauer exit, or if a return to unfavorable conditions could cause the animal to become a full dauer again. We grew *daf-9(dh6)* mutants under unfavorable conditions to induce full dauers, transferred the resulting dauers to favorable conditions to stimulate partial dauer formation, and then transferred them back onto unfavorable conditions to see if they could become full dauers again (**Fig. S1D-F**). Transfer into unfavorable conditions neither dramatically altered pumping rate nor movement speed compared to the mock transfer control, nor did it produce larvae that were similar to full dauers, even 24 hours after a return to unfavorable conditions. This observation suggests that partial dauers may be animals that have committed to, but can only partially complete, dauer exit.

### Evaluating if *daf-9* partial dauers pass through a transient state of full dauer

Under the hypothesis that *daf-9* partial dauers were once full dauers that then partially exited, it should be possible to observe *daf-9* mutants pass through a period of being full dauers before they become partial dauers. We thus grew *daf-9(dh6)* mutants under favorable conditions and scored animals every two hours as being L2d, full dauer, or partial dauer based on important metrics such as pharyngeal pumping, locomotion, and morphology (**Fig. 4A, B,** and see Materials and Methods). Pharyngeal pumping was the most prominent and earliest cue for scoring partial dauers. As a control, we also grew *daf-9(dh6)* and wild-type animals under unfavorable conditions in parallel. To maintain synchrony across the different growth conditions, we grew all animals at a high temperature of 25.5°C but withheld pheromone from the *daf-9(dh6)* mutants grown under favorable conditions. Although growth at 25.5°C favors dauer formation, it alone cannot induce dauer formation in wild-type animals (Ailion and Thomas, 2000). At 44 hours post egg-lay, the vast majority of animals were L2d (**Fig. 4A**). By 49 hours, around 25% of *daf-9(dh6)* mutants grown in the absence of pheromone could be scored as full dauers, while by 52 hours, a surprising 75% of animals were found to be full dauers. After 52 hours, the percentage of full dauers decreased such that by 69 hours, the large majority of animals were partial dauers. In contrast, both the wild-type and *daf-9(dh6)* animals grown under high pheromone conditions showed a steady increase in the proportion of dauers over time, and few if any partial dauers could be found at any time point (**Fig. 4A**). These results show that a proportion of *daf-9(dh6)* mutants grown in the absence of pheromone become full dauers for some period of time.

**Figure 4.**
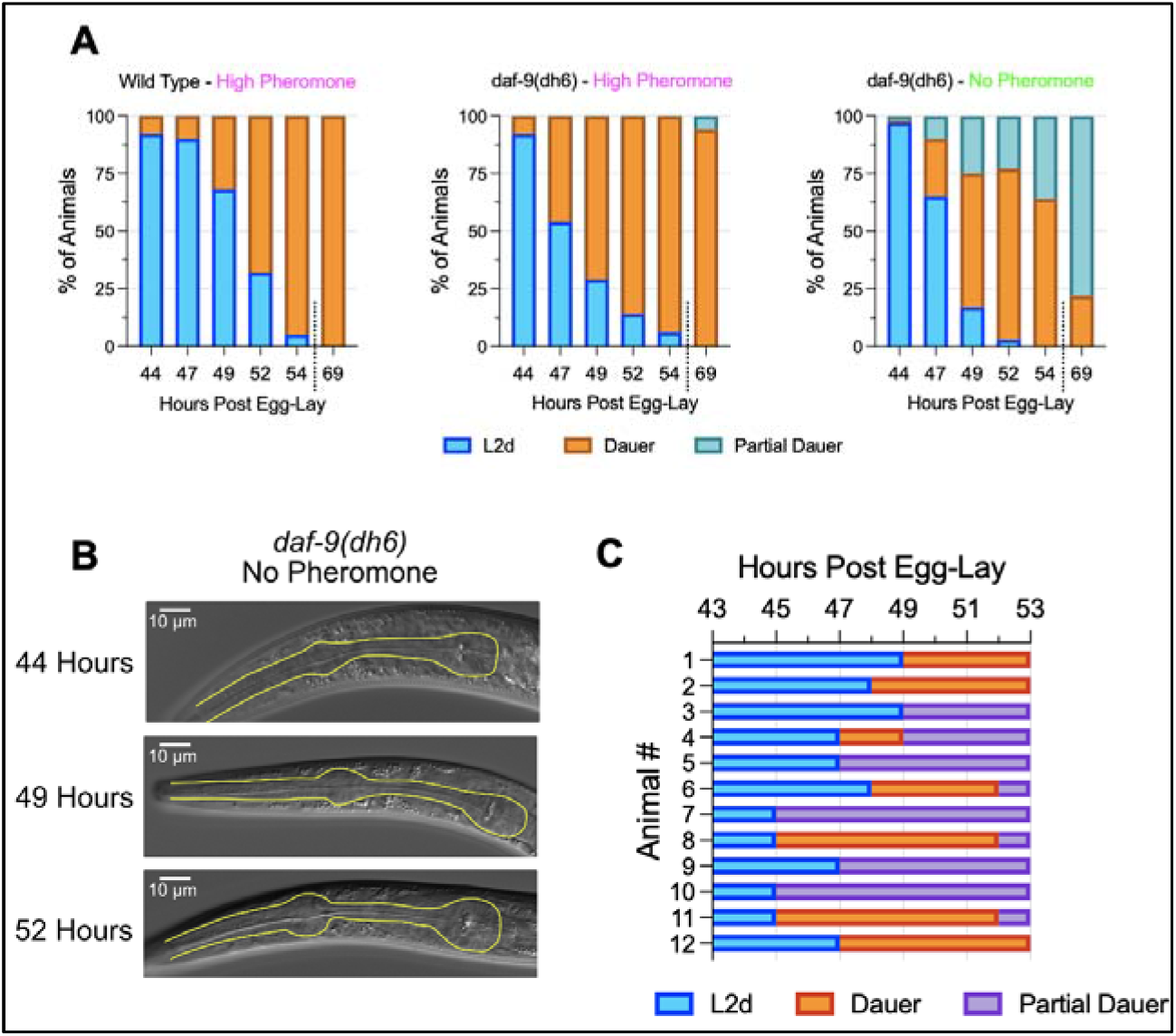
*daf-9(dh6)* larvae transiently become full dauers in the absence of exogenous pheromone. (A) *daf-9(dh6)* mutants were grown in the presence or absence of exogenously added pheromone and then scored as L2d, full dauer, or partially exiting dauer animals as scored by morphology and behavior. n ≥ 53 animals for all observations (B) Representative images of pharynxes from animals grown in conditions identical to the experiment performed in (A). Note the shrinkage of the posterior pharyngeal bulb at 49 hours, indicative of full dauer status, followed by an enlargement of the pharynx at 52 hours, indicative of partial dauer exit. (C) *daf-9(dh6)* mutants grown as in (A), but animals were individually grown and observed every 2 hours starting at 43 hours post egg-lay. 4 of 12 animals could be observed transitioning through a full dauer state before becoming a partial dauer, while other animals (i.e., animal #5) were not observed as full dauers but may have passed the full dauer state in between observation time points.

To determine what fraction of *daf-9(dh6)* mutants pass through a transient dauer state, we repeated the above experiment but with single animals. We grew *daf-9(dh6)* mutants without pheromone at 25.5°C and, after 43 hours post egg-lay, we transferred the resulting L2ds onto new plates without pheromone (one per plate), and we scored individual animals over time (**Fig. 4C**). In concordance with our bulk tracking assay, we observed full dauers between 45 and 50 hours that later became partial dauers. Of 12 tracked animals, we observed seven that went through a period of being full dauers. For these animals, we observed an L2d molt in which the animal detached from and sometimes rolled inside its cuticle (Singh and Sulston, 1978). Afterwards, the animal would cease both movement and pharyngeal pumping before completing radial constriction to become a full dauer. Within a few hours, these dauers slowly began pumping and moving more (a sign of partial dauer exit), but radial expansion did not occur until many hours later (data not shown). Some animals were never observed as having formed full dauers (**Fig. 4C**), which may be because their transition through full dauers occurred in between time points.

We also performed the above single animal observation experiments under more favorable conditions by lowering the temperature to 20°C. However, under these conditions, we were unable to find any *daf-9(dh6)* larvae that went through a full dauer state, despite making observations every hour (**Fig. S2**). Following the same L2d molting process that typically precedes full dauer formation, the animal instead passed through an intermediate state that involved both elements of being a dauer (a darkened body) as well as partial dauer (pumping, motility), before becoming well-recognizable partial dauers usually within one hour. These observations suggest that high temperatures facilitate formation of full dauers in *daf-9(dh6)* mutant animals in the absence of exogenously added pheromone.

### Genetic and physiological factors that could affect partial dauer formation

We sought to characterize the genetic and physiological underpinnings of the *daf-9* partial dauer exit state. We asked whether the *daf-9(dh6)* partial dauer exit phenotype was dependent on other genes in the dauer pathway by performing double mutant analysis of *daf-9(dh6)* with strong loss-of-function mutations in the insulin pathway gene *daf-2(e1370)* (encoding a homolog of the insulin growth factor receptor) and the TGF-β pathway gene *daf-7(e1372)* (encoding an ortholog of human GDF11). Mutants in *daf-2* and *daf-7* possess strong Daf-c phenotypes and form full dauers in the absence of exogenous pheromone at high temperatures. We grew *daf-2(e1370); daf-9(dh6)* and *daf-7(e1372); daf-9(dh6)* double mutants under favorable conditions alongside *daf-9(dh6), daf-2(e1370)*, and *daf-7(e1372)* single mutants to see which would form partial dauers (**Fig. 5A, B).** We found that only *daf-9(dh6)* formed partial dauers under these conditions, while the *daf-2(e1370); daf-9(dh6)* and *daf-7(e1372); daf-9(dh6)* double mutants were phenotypically identical to the *daf-2(e1370)* and *daf-7(e1372)* single mutants in that they formed full dauers. These results indicate that the *daf-9(dh6)* partial dauer phenotype is dependent on the insulin pathway as well as the TGF-β pathway.

**Figure 5.**
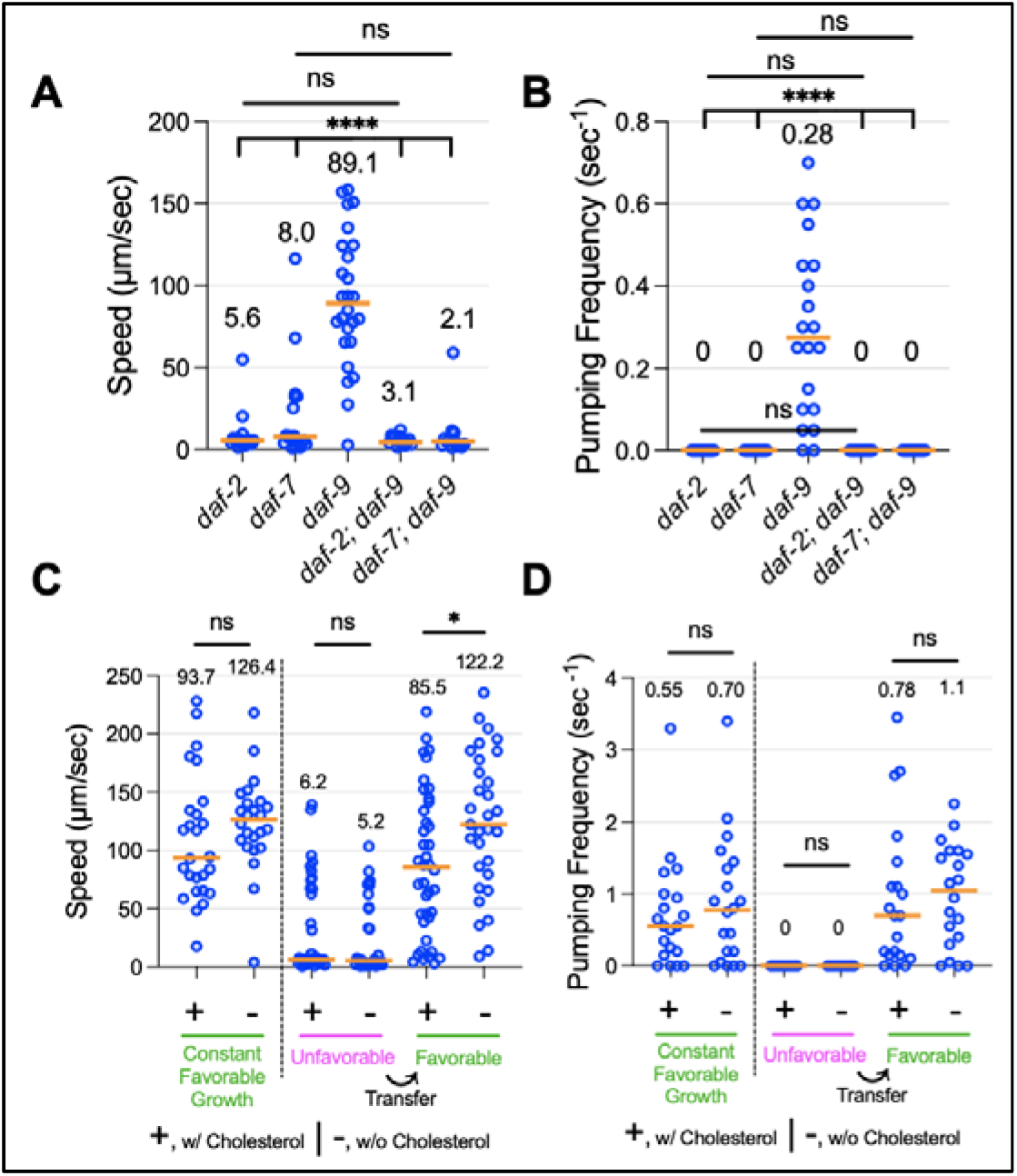
Effects of *daf-2, daf-7*, and cholesterol on the *daf-9* partial dauer phenotype. (A, B) Single and double mutants between *daf-9(dh6)* and the strong loss-of-function mutants *daf-2(e1370)* and *daf-7(e1372)* were grown under favorable conditions. While *daf-9(dh6)* mutants form partial dauers that move faster and pump more frequently, the *daf-2(e1370), daf-7(e1372)*, and their respective *daf-9(dh6)* double mutants do not form partial dauers. p<0.0001 by Mann Whitney test between *daf-9* single mutant and all other double mutants. ns, not significant by Mann Whitney test between double mutants and their single mutant counterparts for *daf-2* or *daf-7*. (C,D) *daf-9(dh6)* partial dauers were formed by constant growth under favorable conditions, or by first inducing full dauers under unfavorable conditions before transferring to favorable conditions to induce dauer exit. In either case, omission of cholesterol from the growth media does not hinder the formation of partial dauers, as scored by locomotion speed and pumping frequency. ns, not significant. *, p<0.05, ****, p<0.0001 by Mann Whitney test. Each dot is one animal, and n ≥ 10 animals for all experimental conditions. Horizontal bars and in-graph numbers show median values.

We speculated that a potential reason for the partial dauer exit phenotype could be that a small amount of reproduction-promoting steroid hormones continues to be produced even in *daf-9* putative null mutants, and that these steroid hormones might trigger partial dauer exit. We reasoned that withholding cholesterol, a precursor for the vast majority of DAF-12 steroid hormone ligands (Aguilaniu et al., 2016), would hinder partial dauer exit formation and keep larvae in their full dauer state. It has been shown that withholding cholesterol exacerbates the developmental defects of many *daf-9* partial loss-of-function mutants by restricting precursor availability (Gerisch et al., 2001; Jeong et al., 2010).

We allowed *daf-9(dh6)* partial dauers to form using two distinct methods: (1) constant growth on favorable conditions and (2) growth under unfavorable conditions followed by a transfer to favorable conditions. We found that withholding cholesterol from the NGM media did not hinder the formation of partial dauers using either of these methods (**Fig. 5C, D**). While these results are inconsistent with the hypothesis that non-*daf-9*-dependent reproduction-promoting steroid hormones promote partial dauer exit, it is also possible that there was sufficient cholesterol or sterol derivatives contained in the medium and/or passed on by previous generations to induce a partial dauer exit state. We attempted to further deprive cholesterol availability by growing *daf-9(dh6)* mutants for two generations in the absence of cholesterol, but these second-generation animals became very sick and could not grow past the L2 stage (data not shown).

### *daf-9* dependent steroid hormone biosynthesis is necessary for and promotes dauer exit

Having confirmed that steroid hormone mutants retain the ability to form full dauers, we proceeded to assess the role of the steroid hormone biosynthesis pathway in dauer exit using *daf-9(dh6)* full dauers for our analyses. *daf-9* encodes a cytochrome P450 enzyme that catalyzes the formation of all known steroid hormones (**Fig. 6A**) (Motola et al., 2006). Among the dafachronic acids, Δ7-DA has been shown to be one of the most effective at rescuing the Daf-c phenotype of *daf-9(dh6)* mutants by allowing them to bypass dauer entry to become healthy adults (Mahanti et al., 2014; Motola et al., 2006). We sought to determine whether Δ7-DA could also rescue the partial dauer exit phenotype of *daf-9(dh6)* larvae (**Fig. 6B**). We let *daf-9(dh6)* full dauers form under unfavorable growth conditions and then transferred them to favorable conditions with varying concentrations of Δ7-DA and scored for complete dauer exit the next day. At low Δ7-DA concentrations, animals become partial dauers, while 100 nM Δ7-DA is sufficient to fully rescue the dauer exit phenotype of *daf-9(dh6)* mutants by prompting nearly all animals to exit the dauer state and resume reproductive development. Nonlinear regression analysis of the dose response curve reveals an EC_50_ of 7.56 nM. We also determined whether Δ7-DA could induce dauer exit of *daf-9(dh6)* mutants in the presence of pheromone. Even at 100 nM Δ7-DA, all animals remained as full dauers (data not shown), which could be because their lack of feeding and/or their thickened cuticle (Cassada and Russell, 1975) preclude access to Δ7-DA.

**Figure 6.**
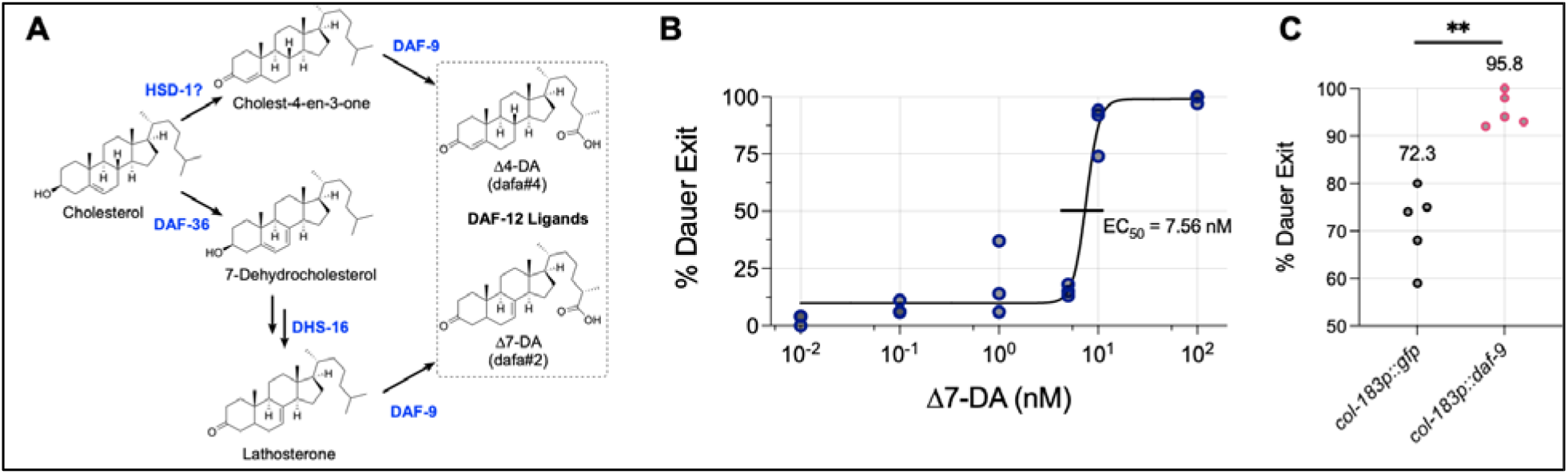
The *daf-9* steroid hormone pathway is necessary for and promotes dauer exit. (A) Steroid hormone biosynthetic pathway. DAF-9 catalyzes the final oxidation step required to form all known endogenous ligands of DAF-12, which promote reproductive development over dauer. Other ligands not shown include Δ0-DA and Δ1,7-DA, whose biosynthetic pathways are not well characterized. Figure adapted from Mahanti *et al*. 2014 and Aguilaniu *et al*. 2016. (B) Dose response curve for the efficacy of Δ7-Dafachronic Acid (Δ7-DA) to rescue the dauer exit defect of *daf-9(dh6)* mutants. *daf-9(dh6)* animals were induced to form full dauers under unfavorable conditions, and then transferred to favorable conditions containing various concentrations of Δ7-DA. 24 hours later, animals were scored for complete dauer exit as determined by substantial radial expansion and lengthening. Lower concentrations yielded partially exited dauers, while 100 nM induced nearly all animals to completely exit dauer. (C) Overexpression of *daf-9* from the hypoderm and dauer-specific promoter *col-183p* promotes dauer exit. For (B) and (C), each dot is a biological replicate of scored dauer exit plates, with each group having a total n ≥ 100 animals. **, p<0.01 by bootstrapped permutation test using 10^4 samples.

We next determined whether overexpression of *daf-9* could be sufficient to induce dauer exit even in a wild-type background. Since constitutive and ubiquitous overexpression of *daf-9* would likely preclude dauer formation, we drove expression of *daf-9* cDNA from the *col-183* promoter (Shih et al., 2019). *col-183* shows maximal expression during dauer and within the hypoderm, a tissue that exhibits high *daf-9* expression levels during reproductive development (Gerisch et al., 2001; Schaedel et al., 2012). Overexpression of *daf-9* from the *col-183* promoter significantly increased the fraction of dauers that exited, indicating that hypodermal *daf-9* expression during dauer can promote dauer exit (**Fig. 6C**).

### Spatiotemporal regulation of *daf-9* during dauer exit resembles that of dauer entry

We characterized the spatiotemporal regulation of *daf-9* during the dauer exit process to examine whether it differs from that during dauer entry. Prior to the dauer entry decision, *daf-9* is expressed exclusively in the XXX neuroendocrine cells. When animals decide to enter the reproductive life cycle, *daf-9* expression increases throughout the hypodermis (Gerisch et al., 2001; Schaedel et al., 2012). To test whether this expression pattern holds true during dauer exit, we used the same DAF-9::GFP translational fusion-bearing strain (Gerisch et al., 2001) and monitored GFP expression as the animals exited dauer (**Fig. 7A,B**). Upon shifting dauer larvae from unfavorable to favorable conditions to induce exit, we observed an increase in the proportion of animals displaying hypodermal GFP expression. The proportion peaked at 18 hours post-shift, which is when larvae have nearly entered L4, at which point nearly 75% of animals showed hypodermal GFP. After 36 hours post-shift, the vast majority of animals (~90%) lost all hypodermal GFP but retained GFP expression in the XXX cells. Thus, the spatiotemporal dynamics of *daf-9* expression for the dauer exit decision seem to match those of the dauer entry decision in that there is widespread hypodermal *daf-9* expression as the animal chooses the reproductive route during development.

**Figure 7.**
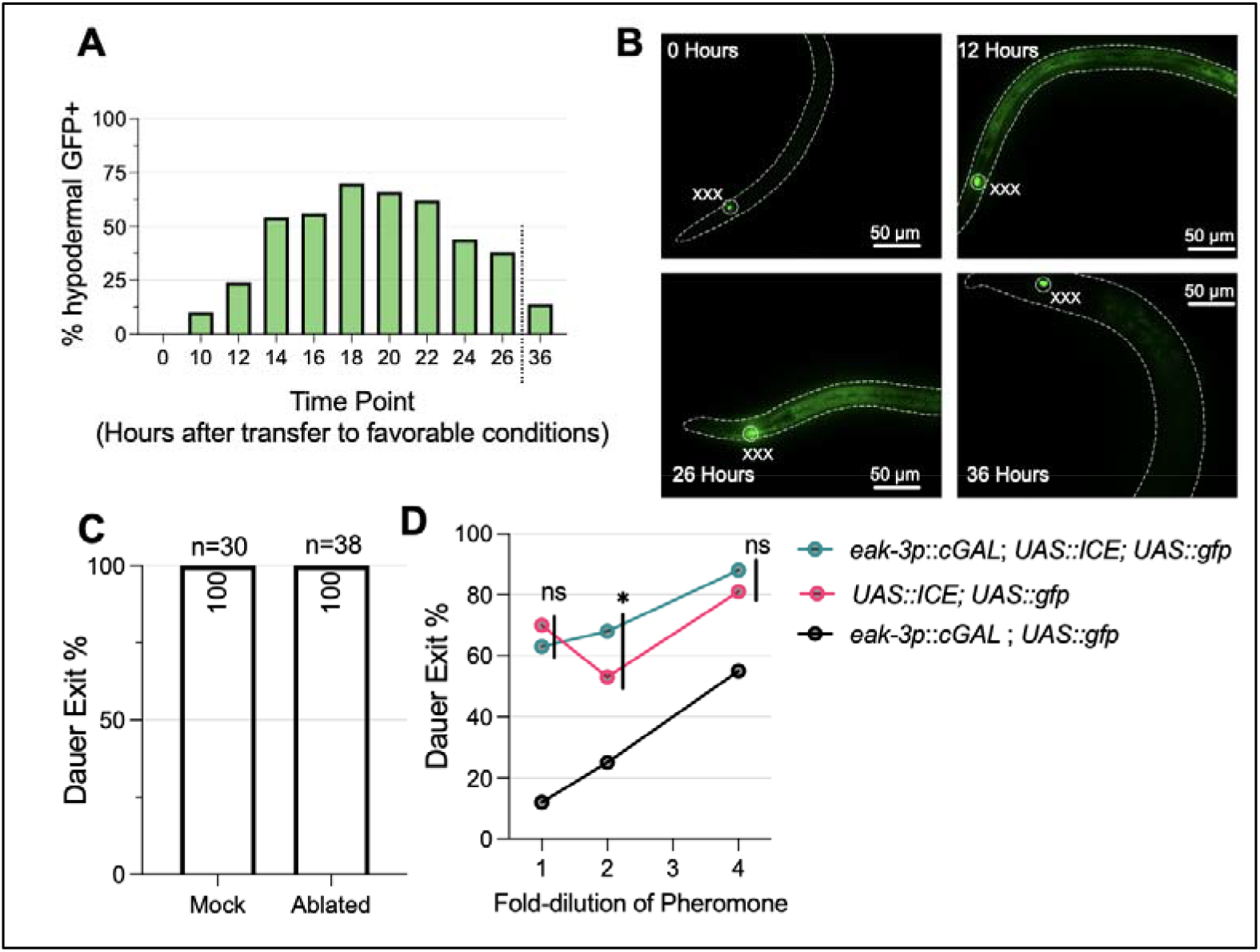
Spatiotemporal dynamics of *daf-9* expression during dauer exit. (A) Time course of hypodermal *daf-9::gfp* expression during dauer exit. Dauers expressing a *daf-9::gfp* translational fusion were transferred to favorable conditions and imaged for the presence of absence of GFP signal in the hypoderm. *daf-9::gfp* is expressed in the highest percentage of animals at ~16-20 hours after transfer to favorable conditions. n ≥ 20 for all time points. (B) Representative images of animals observed in (A), with bodies outlined and the XXX neuroendocrine cells (which constitutively express *daf-9::gfp*) labeled. Note the increase in DAF-9::GFP in images for 12 and 26 hours. (C) Laser ablation of XXX cells does not prevent dauer exit. Following ablation of the XXX cells using a laser microbeam, dauers and mock-ablated control dauers were transferred to favorable conditions and scored for dauer exit 24 hours later. (D) Genetic ablation of the XXX cells does not drastically impact dauer exit. Dauers whose XXX cells were genetically ablated were transferred to plates with decreasing pheromone concentrations and scored for dauer exit 24 hours later. n ≥ 102 animals for each sample. ns, not significant. *, p<0.05 by bootstrapped permutation test for a proportion using 10^4 samples.

The constitutive expression of *daf-9* in the XXX cells throughout dauer led us to ask whether the XXX cells might be important for dauer exit. Published reports indicate if the XXX cells are ablated by a laser microbeam during the L1 stage, only a small fraction of animals become dauer-like larvae even under favorable conditions (Gerisch et al., 2001; Ohkura et al., 2003). However, if ablation occurs during the L2d stage after the animals have been reared under unfavorable conditions, then nearly all larvae form dauers even after a shift to favorable conditions (Schaedel et al., 2012). To examine whether the XXX cells are dispensable for dauer exit, we bilaterally ablated the XXX cells in dauers using a laser microbeam in dauer animals and transferred the ablated animals to a recovery plate under favorable conditions to induce dauer exit. We found that all XXX-ablated dauer larvae were able to exit dauer, similar to their mock ablated counterparts (**Fig. 7C)**. To validate this finding, we also genetically ablated the XXX cells by expressing the human caspase gene *ICE* from the XXX-specific promoter *eak-3p* using our cGal bipartite expression system for *C. elegans* (Wang et al., 2017). We confirmed ablation of the XXX cells by noting the loss of GFP in XXX under fluorescence microscopy in a strain expressing the *UAS::ICE* construct (**Fig. S3**). With the enhanced throughout of the genetic ablation method compared to laser ablation, we recovered animals onto different pheromone concentrations to tease out more subtle changes in dauer exit phenotypes. Genetic ablation of the XXX cells did not substantially prevent dauer larvae from exiting more when compared to control animals expressing the *UAS::ICE* effector transgene without the XXX cell-specific driver (**Fig. 7D**). In tandem with our laser ablation results, these findings suggest that the XXX cells may not be essential for dauer exit.

## Discussion

### Partial dauers formed by steroid hormone mutants are likely partially exited dauers

In this study, we evaluated how the steroid hormone pathway regulates both the dauer entry and dauer exit developmental decisions by first addressing why *daf-9* mutants form partial dauers. Our evidence favors the hypothesis that *daf-9* partial dauers, along with other partial dauers formed by mutations in steroid hormone biosynthesis, represent a state in which dauers have commenced but cannot complete dauer exit. We find that forming *daf-9* full dauers under unfavorable conditions followed by transfer to favorable conditions to induce dauer exit produces animals that resemble the partial dauers that are formed when *daf-9* mutants are grown constantly under favorable conditions (**Fig. 2, 3E-G**). We also show that even when pheromone is omitted from the growth medium, some *daf-9(dh6)* animals pass through a state of full dauer before initiating dauer exit owing to the favorable environment (**Fig. 4**). But given the lack of reproduction-promoting steroid hormones such as Δ7-DA, these larvae can only partially exit from dauer, resulting in a partial dauer state that slowly grows to unhealthy and sterile adulthood.

Under completely favorable conditions (i.e., no pheromone and low temperature), *daf-9(dh6)* larvae could not be found in a full dauer state (**Fig. S2**). Following the L2d molt, we were only able to find *daf-9(dh6)* mutants in a transient, indeterminate state that looked like a hybrid between an L2d and a partial dauer in terms of morphology and behavior. Within an hour, these animals then quickly went on to become familiar partial dauers. While it is difficult to speculate why this intermediate state arises, one possibility has to do with the fact that dauer formation affects multiple tissues, including the epithelium, intestine, pharynx, and body muscles (Androwski et al., 2017). While this process normally happens synchronously, in the case of *daf-9(dh6)* mutants grown under favorable conditions, there could be asynchrony as different tissues start to engage dauer exit programs before other tissues finish executing dauer entry programs. For instance, as *daf-9(dh6)* mutants conclude the L2d molt, the nervous system could immediately sense food and the lack of pheromone to trigger dauer exit programs that promote pumping and locomotion. These dauer exit behaviors could start before epidermal tissues finish the radial constriction process associated with dauer entry. The result would be an animal that looks and behaves neither like a partially exited dauer nor a full dauer, as we see in our experiments. When conditions are tuned to be slightly less favorable (such as by increasing the temperature), the different tissues of *daf-9* mutants can engage dauer entry in a more synchronous fashion, therefore allowing observation of the full dauer state by eye. Another possibility to explain these results is simply that *daf-9* mutants completely skip or fail to enter the full dauer state under completely favorable growth conditions because some tissues never fully execute their dauer entry programs, but why that might happen remains unclear. Experiments that can track how individual tissues engage dauer entry and exit programs will be helpful in teasing these possibilities apart.

Our conclusions regarding the *daf-9* partial dauer state as being one of partial dauer exit aligns with those of a previous group concerning the partial dauer phenotype of *daf-12* Daf-c alleles such as *daf-12(rh273)*, in which they suggest that *daf-12* Daf-c mutants initiate all the programs necessary for full dauer entry, but immediately detect favorable conditions upon dauer entry and thus promptly attempt to exit (Antebi et al., 1998). In further support of these conclusions, we find that the Daf-c *daf-12(rh273)* strain phenocopies *daf-9(dh6)* when it comes to partial dauer exit (**Fig. S1B, C**).

### The *daf-9* partial dauer state provides new insights into the dauer exit process

Recognizing that the *daf-9* partial dauer state may be a partially exited dauer raises many intriguing questions about dauer exit biology. It prompts a consideration of what genetic, developmental, and physiological factors may be responsible for triggering partial dauer exit. We considered that residual steroid hormone production, either through non-*daf-9*-dependent biosynthesis or transgenerational rescue, suffices to trigger partial dauer exit. The former is improbable because there are no characterized biochemical pathways to form DAF-12 ligands that do not involve DAF-9 (Aguilaniu et al., 2016). The latter, while difficult to prove since we could not limit cholesterol deprivation past one generation, seems unlikely. Growing *daf-9(dh6)* mutants on media lacking cholesterol does not suppress partial dauer formation (**Fig. 5C, D**). Moreover, a double mutant for both *daf-9* and *daf-36*, which would presumably have lower steroid hormone levels than *daf-9* single mutants alone, phenocopies *daf-9* to produce partial dauers (Rottiers et al., 2006).

The partial dauer exit phenotype can be suppressed by mutations in important components of the dauer pathway such as DAF-2/InsR and DAF-7/GDF11 (**Fig. 5A, B**). This is consistent with a model in which steroid hormone mutants form partial dauers because insulin and TGF-β pathways are activated in response to the animal sensing favorable conditions, subsequently triggering initial dauer exit behaviors and morphologies. This model is further supported by the fact that the transcriptional outputs of the insulin pathway, mediated by DAF-16/FOXO, are distinct from those of DAF-12/FXR, although the two pathways have significant crosstalk (Jeong et al., 2010). While DAF-16/FOXO directly regulates the transcription of genes related to metabolism, stress resistance, and longevity, DAF-12/FXR is known to transcriptionally regulate heterochronic gene pathways that govern developmental timing (Hochbaum et al., 2011; Kumar et al., 2011). Given these different transcriptional networks, one hypothesis is that favorable conditions activate insulin and TGF-β pathways that transcriptionally upregulate target genes to initiate dauer exit. However, without liganded DAF-12/FXR activity to stimulate heterochronic gene expression, the larvae cannot proceed to reproductive adulthood. Further experiments that precisely manipulate insulin, TGF-β, and steroid hormone pathway activity in full and partial dauers will help delineate such a model. The *daf-9* partial dauer exit state, therefore, provides an avenue by which other regulatory components underlying dauer exit can be explored.

Could other described partial dauers, such as those produced by double mutant strains carrying *daf-16* (Ailion and Thomas, 2000; Vowels and Thomas, 1992), also be characterized as partially exited dauers? The features of those partial dauers closely match those of the *daf-9* partial dauers that we and others have described including pumping, an enlarged pharynx, and intermediate intestinal darkness (Gerisch et al., 2001; Vowels and Thomas, 1992). One distinction is that these double mutant partial dauers were described to quickly and spontaneously exit to adulthood (Vowels and Thomas, 1992), while *daf-9(dh6)* partial dauers cannot ever fully exit. The spontaneity of complete exit in the double mutant partial dauers suggests that they may be in a partially exited dauer state waiting for a threshold event to complete dauer exit. We have not rigorously tested those strains using our partial dauer exit analyses, and it remains to be determined whether these double mutant partial dauers have the capacity to enter full dauers under unfavorable conditions as can steroid hormone mutants.

### Steroid hormone biosynthesis governs dauer exit in a manner similar to the L1 to L2 versus L2d dauer entry decision

We evaluated how the *C. elegans* steroid hormone pathway regulates dauer exit in comparison to dauer entry. We established that steroid hormones are essential for full dauer exit by showing that *daf-9(dh6)* dauers become competent for complete dauer via supplementation of Δ7-DA at an EC50 in the low nanomolar range (**Fig. 6B**). Previous dose response curves showing the relationship between Δ7-DA and the Daf-c dauer entry phenotype of *daf-9(dh6)* animals show an EC50 of ~5-25 nM, while 100 nM fully rescues the dauer entry phenotype (Schaedel et al., 2012). These comparable results suggest that similar concentrations of Δ7-DA mediate both dauer entry and dauer exit in *daf-9(dh6)* animals. We further show that hypodermal overexpression of *daf-9* during dauer promotes dauer exit **(Fig. 6C**), therefore demonstrating a parallel role for *daf-9*-dependent steroid hormones in regulating both dauer entry and dauer exit.

The dauer entry decision can be split into two distinct subdecisions, both of which are regulated by the steroid hormone pathway: (1) the L1 to L2 versus L2d and (2) the L2d to L3 versus dauer (Golden and Riddle, 1984). We asked whether the steroid hormone pathway regulates the dauer exit decision in a manner that resembles the former, the latter, or neither dauer entry subdecision. Our results collectively suggest that the role of the steroid hormone pathway in dauer exit more closely mirrors that of the L1 to L2 versus L2d decision for multiple reasons.

First, our *daf-9::gfp* translational fusion analysis shows that hypodermal upregulation of *daf-9::gfp* begins around 10-14 hours following transfer of dauers onto favorable conditions (**Fig. 7A**). This delay in hypodermal *daf-9::gfp* expression nearly matches that of the L1 to L2 vs. L2d decision, in which it was shown that *daf-9::gfp* fluorescence increased starting at 24 to 27 hours post hatch in animals grown under favorable conditions, while larvae commit to reproductive adulthood much earlier at around 14 to 16 hours post hatch (Schaedel et al., 2012). In stark contrast, during the L2d to dauer versus L3 decision, hypodermal DAF-9::GFP fluorescence closely aligned with the time window in which L2d larvae committed to reproductive adulthood (Schaedel et al., 2012). Importantly, given that dauer exit commitment occurs within one to two hours following transfer onto favorable conditions (Golden and Riddle, 1984), the fact that hypodermal DAF-9::GFP fluorescence does not appear until hours later suggests that hypodermal upregulation of *daf-9* may be a consequence of, rather than a cause of, the commitment to exit dauer.

Secondly, our XXX ablation experiments argue against an essential role for the XXX cells in dauer exit, as laser or genetic ablation of XXX cells do not prevent dauers from exiting (**Fig. 7C, D)**. Such observations are consistent with the nonessential role of XXX cells in the L1 to L2 versus L2d decision, in which groups have reported that laser ablation of XXX cells in L1 larvae has only a minor effect on dauer entry (Gerisch et al., 2001; Ohkura et al., 2003). In contrast, the XXX cells are required for reproductive adulthood in the L2d to L3 versus dauer subdecision, as ablation of the XXX cells in L2d larvae prevents development into the L3 stage even under favorable conditions (Schaedel et al., 2012). Taken together, these results suggest higher similarity between dauer exit and the L1 to L2 versus L2d dauer entry decision, at least as far as the *daf-9*-dependent steroid hormone pathway is concerned.

### Dauer entry and dauer exit are asymmetric developmental decisions

Despite dauer entry and dauer exit sharing similar features, including their sensory inputs and regulatory pathways, our work highlights that these two decisions are inherently asymmetric. One basic distinction is that dauer entry comprises two developmental subdecisions, while dauer exit consists of just one. Moreover, the two dauer entry subdecisions involve a choice between two mutually exclusive developmental paths that must be chosen within a certain timeframe. On the other hand, dauer exit has a different decision architecture involving a continuous decision made over an indeterminate timeframe (i.e., animals will exit dauer or they will stay as dauers). The differences in these decision-making structures are highlighted by the *daf-9* partial dauer exit phenotype in that there does not seem to be an analogous “partial dauer entry” phenotype for *daf-9* mutants. This difference is because the dauer entry decision does not allow an animal to arrest while deciding, but instead forces the animal to commit to one of the two developmental pathways once enough time has elapsed. Such examples of asymmetry in diapause-related developmental decision-making yield important implications for how the entry and exit decisions might be differentially regulated.

### Hormonal regulation of diapause entry and exit in other organisms

The DAF-12 steroid hormone pathway in *C. elegans* is conserved in other parasitic nematode species, whose infective larvae stage are comparable to *C. elegans* dauers. Treatment of multiple parasitic species during early larval growth with Δ7-DA prevents entry into the infective stage, while treatment of infective larvae prompts exit from the infective stage (Ogawa et al., 2009; Wang et al., 2009), mirroring our results using Δ7-DA to induce full dauer exit (**Fig. 6B**). Such conversation suggests that mechanistic knowledge of how steroid hormones control dauer exit in *C. elegans* could yield potential therapeutic insights to combat other parasitic species.

Diapause is evolutionarily conserved and phylogenetically widespread, and hormonal control of both diapause entry and exit is especially well-studied in insects (Denlinger et al., 2012). In *Heliothis* and *Helicoverpa* species of moth, diapause entry is likely caused by insufficient levels of diapause hormone (DH) and prothoracicotropic hormone (PTTH) (Xu and Denlinger, 2003). Administration of DH or DH mimics to diapausing pupae results in diapause termination, indicating that DH is sufficient to cause diapause exit (Zhang et al., 2011). The notion that diapause occurs in the absence of a pro-development hormone(s) and that diapause terminates in its presence may therefore be conserved between nematodes and insects, but in some insect species where diapause is maternally controlled, diapause entry and exit are regulated via different hormonal processes. Studies of embryonic diapause in the silkworm *Bombyx mori* have demonstrated a central role for DH in triggering, rather than preventing, diapause in developing embryos (Sato et al., 1992). Diapause termination, on the other hand, does not appear to depend on the absence of DH but instead on the presence of pro-development ecdysteroids such as 20-hydroxyecdysone (Iwata et al., 2005; Sonobe and Yamada, 2004). In *C. elegans* and other animal species, co-opting the same hormonal signaling process for preventing diapause entry and inducing diapause exit could be an adaptive strategy that efficiently uses pre-existing molecular pathways for multiple purposes.

## Materials and Methods

### *C. elegans* strains and maintenance

*C. elegans* strains were derived from the wild-type strain N2 (Bristol) and were cultured according to standard laboratory conditions on Nematode Growth Medium (NGM) agar seeded with *E. coli* OP50 as the food source. A list of strains used in this study, including their genotypes and origins, can be found in Table S1. Maintenance and propagation of *C. elegans*, with the exception of *daf-9* loss-of-function mutants, were performed under typical growth conditions with Nematode Growth Medium (NGM) agar seeded with OP50 *E. coli* cultures as described previously (Brenner, 1974). The *daf-9(dh6), daf-9(e1406)*, and *daf-9(m540)* mutants were propagated in the presence of 100 nM Δ7-Dafachronic Acid (Δ7-DA) to promote reproductive development. The *daf-9(dh6)* strain was obtained by propagating non-array carrying animals from PS5511 (*daf-9(dh6); dhEx24[T13C5, pTG96(+)]*) on Δ7-DA.

### Dauer Entry Induction

To induce full dauers in wild-type and *daf-9* mutants, 10-20 young adults were placed on 35 mm diameter Petri dishes containing 2 mL of NGM agar (without peptone) supplemented with a quantity of crude pheromone extract (Schroeder and Flatt, 2014) that normally induce 95-100% of dauers in wild-type animals – typically 10-30 μL per 2 mL of agar. Plates were seeded with 10 μL of 8% w/v *E. coli* OP50 that was heat-killed at 95°C for 5 minutes. Adults were picked onto the plate and allowed to lay eggs at room temperature (RT; 22-23°C) for 5-9 hours before being removed, during which time they typically laid 200-300 eggs. The plates were then further seeded with an additional 20 μL of heat-killed OP50. Afterwards, the plates were wrapped with parafilm and incubated at 25.5°C for 60-72 hours.

### Dauer Exit Assay

Dauers were formed according to Dauer Entry Induction, above. In most cases, dauers were selected for by an SDS wash (2%, 30 minutes, 25°C) to kill non-dauers before being washed 3x in M9 solution (1 minute, room temperature, 1000 x *g*). Surviving dauers were then plated onto dauer exit plates, which were identical to dauer entry plates but contained a lower concentration of crude pheromone extract (typically 0.5-2 μL per 2 mL of agar) that had been determined to induce ~40-60% of wild-type dauers to exit within 24 hours. In the case of the genetic ablation assay using the *UAS::ICE* construct, the SDS wash step was omitted since dauers bearing the *UAS::ICE* construct were SDS sensitive. Dauers were instead washed directly onto the dauer exit plate. Following 24 hours after dauers were transferred onto dauer exit plates, dauer exit was scored using the following criteria. Larvae were scored as having exited dauer if they showed any pharyngeal pumping or if their body had thickened and lightened considerably. Additional factors that favored scoring an animal as having exited dauer included whether the larva showed foraging behavior (such as increased head turns) or increased and consistent locomotory behavior, both of which are absent from dauers.

### Microscopy and Image Analysis

Worms were immobilized on a 4-10% agarose pad (10% for dauer imaging, 4% for others) in 1-2 μL of 10 mg/mL levamisole or 50 mM sodium azide. Imaging was performed on a Zeiss AxioImager2 equipped with a Colibri 7 for LED fluorescence illumination and an Axiocam 506 Mono camera (Carl Zeiss Inc., White Plains, NY). Pharyngeal bulb width measurements were performed using Zen Blue 2.3 (Zeiss) software by using the length tool to measure the widest section of the posterior pharyngeal bulb. Images were processed using FIJI (ImageJ).

### Laser Ablation in dauers

PS8568 animals expressing *gfp* in the XXX cells were induced to form dauers and then immobilized on a 4% agarose pad with 10 mg/mL levamisole. Laser ablation was performed on a Zeiss Axioskop (Carl Zeiss Inc.) equipped with an Andor MicroPoint nitrogen-pulsed laser microbeam (Oxford Instruments, Abingdon, UK). XXX cells were visualized under fluorescence, and the laser was fired at ~5 Hz for a total 20-30 pulses, or until all discernable fluorescence was gone. Cellular damage could often be visualized under DIC. Following surgery, animals were recovered onto a 35 mm NGM plate seeded with OP50 washed in S Basal and scored for dauer exit 24 hours later. GFP was no longer discernable under stereomicroscopy in successfully ablated animals. Mock ablated animals were prepared and rescued identically to ablated animals but without receiving laser treatment.

### Partial Dauer Induction

Partial dauers of *daf-9* mutants were successfully obtained using two methods. The first method (favorable growth) is as follows. 10-20 young adults were placed on 35 mm Petri dishes containing 2 mL of NGM agar (without peptone) seeded with 10 μL of 8% w/v OP50 washed twice in S Basal and including 50 μg/mL streptomycin to limit bacterial growth (Golden and Riddle, 1984). Adults were allowed to lay eggs at RT before being removed, after which an additional 20 μL of 8% S Basal-washed OP50 was added. Plates were wrapped in parafilm and grown at 20°C for 60-72 hours to yield partial dauers. The second method (unfavorable growth followed by transfer to favorable conditions) involved forming dauers according to Dauer Entry Induction. Dauers were then washed in 2% SDS (30 minutes, 25°C) before being washed 3x in M9, collected by centrifugation (1 minute, room temperature, 1000 x *g*), and then plated onto 35 mm NGM Petri plates lacking pheromone. Partial dauers could be found 24 hours later.

### Behavioral Scoring

For all behavioral scoring events, animals were first allowed to acclimate to room temperature for thirty minutes prior to scoring. For pumping frequency scoring, animals were manually scored under a stereomicroscope at 100x magnification over a twenty second period. A pumping event was scored as a contraction of the pharyngeal grinder. For locomotion analysis, one-minute videos were recorded and analyzed using the WormLab Imaging System and software (MBF Bioscience, Willston, VT). Videos contained on average 8-12 animals per recording event, and multiple videos were pooled together for each experiment. Recordings were performed in areas of the plate that were away from food to maximize contrast, since recordings performed on food prevented accurate tracking of Animals. In all cases, the Peristaltic Speed (Absolute) output, measured in μm/second, was reported for these experiments. We note that the non-zero absolute speed output of many plotted dauers (i.e., Figure 3) were a result of noise owing to unstable camera movements. These dauers were often perfectly still when viewed by eye (see Video 1).

### Time course tracking of dauer status in *daf-9(dh6)* mutants

Wild-type and *daf-9(dh6)* full dauer controls were grown according to Dauer Entry Induction. Simultaneously, *daf-9(dh6)* mutants were grown in the absence of exogenous pheromone. All strains were grown at 25.5°C in order to maximize developmental synchrony across the different conditions. For single animal tracking of *daf-9(dh6)* partial dauers, individual L2ds grown under the conditions described above were picked onto new non-pheromone-containing plates starting at 43 hours post egg-lay and scored every two hours. An animal was scored as being a partial dauer if any pumping was observed and/or the body thickened and lightened compared to a normal dauer.

### Cholesterol Deprivation

Cholesterol deprived plates were made similarly to normal dauer plates (i.e., in 35 mm petri dishes with 2 mL NGM agar without peptone) except that ethanol solvent was added in place of cholesterol. This method was sufficient to enhance the Daf-c phenotype of *daf-9(m540)* mutants, as previously described (Jeong et al., 2010). More severe methods of cholesterol starvation, such as using agarose in place of agar and passaging the animals over two generations in the absence of cholesterol (Gerisch et al., 2001), yielded unhealthy larvae that could not grow to become dauers.

### Double mutant analysis between *daf-9(dh6)* and *daf-2(e1370)* or *daf-7(e1372)*

Double mutants were constructed as follows. Wild-type males were mated to the balancer strain *sC1[dpy-1(s2170) umnIs21]*, which are marked by a recessive Dumpy (Dpy) phenotype and a dominant pharyngeal GFP phenotype. The resulting male sC1heterozygotes were mated to *daf-9(dh6); dhEx24* hermaphrodites. *dhEx24* is an extrachromosomal array containing the cosmid T13C5, containing a rescuing wild-type copy of the *daf-9* locus and a *sur-5p::gfp* marker expressing GFP throughout the body. Hemizygous *daf-9(dh6)*/0 male progeny with both whole-body GFP+ (from *dhEx24*) and pharyngeal GFP+ (from *sC1*) were mated again to *daf-9(dh6) dhEx24* hermaphrodites, and the Dpy F_2_ double GFP+ progeny were obtained to yield *sC1(s2023) [dpy-1(s2170) umnIs21]; daf-9(dh6); dhEx24*. This strain was then crossed into a *daf-2(e1370)* or *daf-7(e1372)* background to yield *sC1(s2023) [dpy-1(s2170) umnIs21]; daf-9(dh6); dhEx24*. The *sC1* balancer could then be used to follow *daf-2(e1370)* and *daf-7(e1372)* in *trans* to facilitate the construction of double mutatns between these mutations and *daf-9(dh6)*. For pharyngeal pumping and locomotion assays, the balanced double mutants were grown from eggs at 25.5°without pheromone for three days, and *daf-2(e1370); daf-9(dh6)* or *daf-7(e1372); daf-9(dh6)* dauer larvae were identified by looking for non-GFP dauers and picked onto new plates. These dauers were allowed to acclimate on the new plates for at least thirty minutes prior to scoring.

### Statistical Analysis and Plotting

Plots were designed using Prism 9.0 (GraphPad, San Diego, CA). Dose response curves for steroid hormones were calculated using Prism’s nonlinear regression tool ([Agonist] vs response) with four parameters and the EC50 parameter constrained to be greater than zero. Mann-Whitney tests were performed in Prism. Permutation tests for dauer exit proportion between two samples were performed by first binarizing dauer exit data, pooling the two samples together, and simulating experiments by drawing two samples out of the pooled binarized data. The p value was calculated by comparing the number of simulated experiments, out of 10^4^, in which the difference between the simulated proportions was greater than the observed difference between the actual proportions.

## Acknowledgements

Δ7-DA was a gift from the lab of Frank Schroeder (Cornell University). We thank Mengyi Cao and Stephanie Nava for the XXX cGAL driver. Some strains were provided by the CGC, which is funded by NIH Office of Research Infrastructure Programs (P40 OD010440). We are grateful to members of the Sternberg lab for their feedback on the manuscript, particularly Hillel Schwartz.

## Author Contributions

M.G.Z. and P.W.S. conceived of the study. M.G.Z. performed the experiments and analyzed the data. M.G.Z. wrote the manuscript with editorial assistance from P.W.S.

---

**Video 1:**
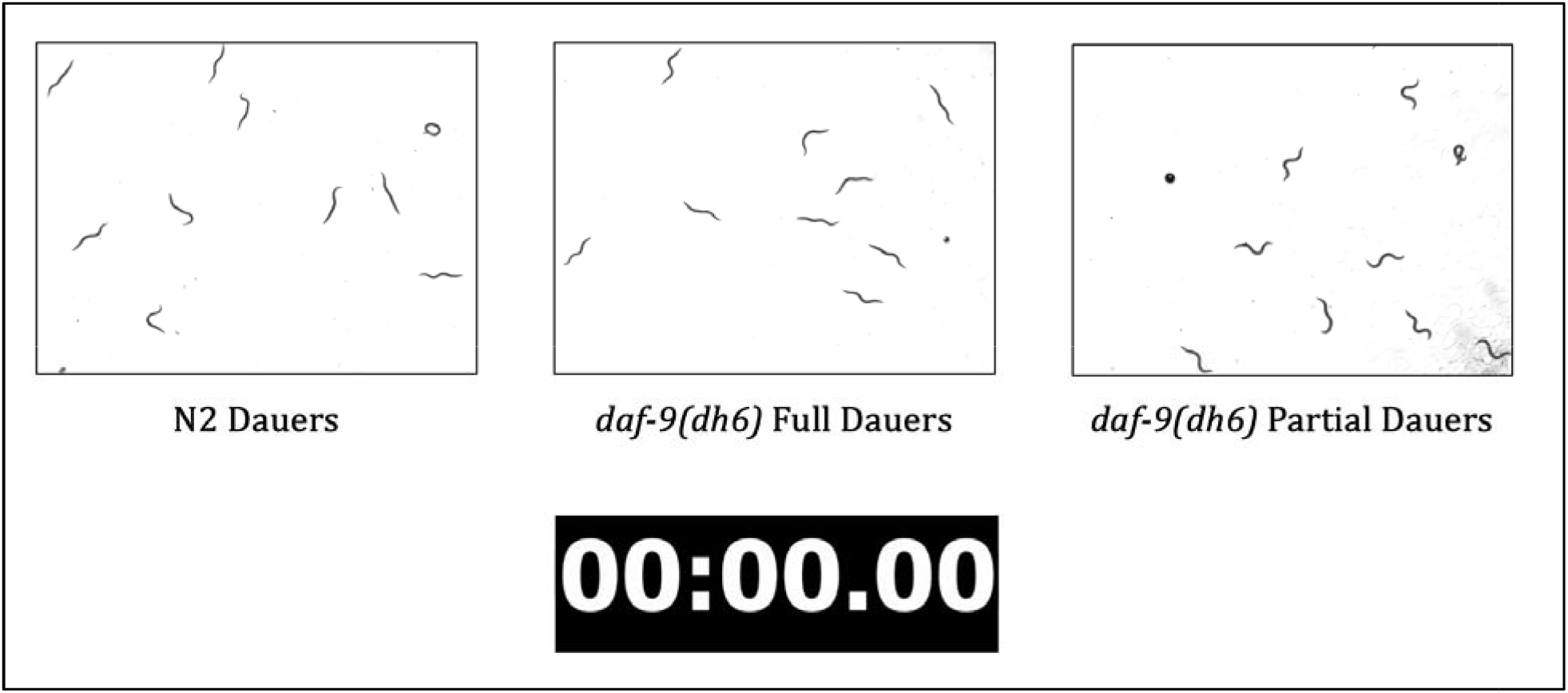
*daf-9(dh6)* full dauers resemble wild-type dauers in terms of locomotion. Shown are 1-minute video recordings of N2 (wild-type) and *daf-9(dh6)* dauers formed under unfavorable growth conditions in comparison to *daf-9(dh6)* partial dauers formed under favorable growth conditions. N2 (wild-type) and *daf-9(dh6)* full dauers move mu**ch** more slowly, if at all, compared to *daf-9(dh6)* partial dauers.

**Video 2:**
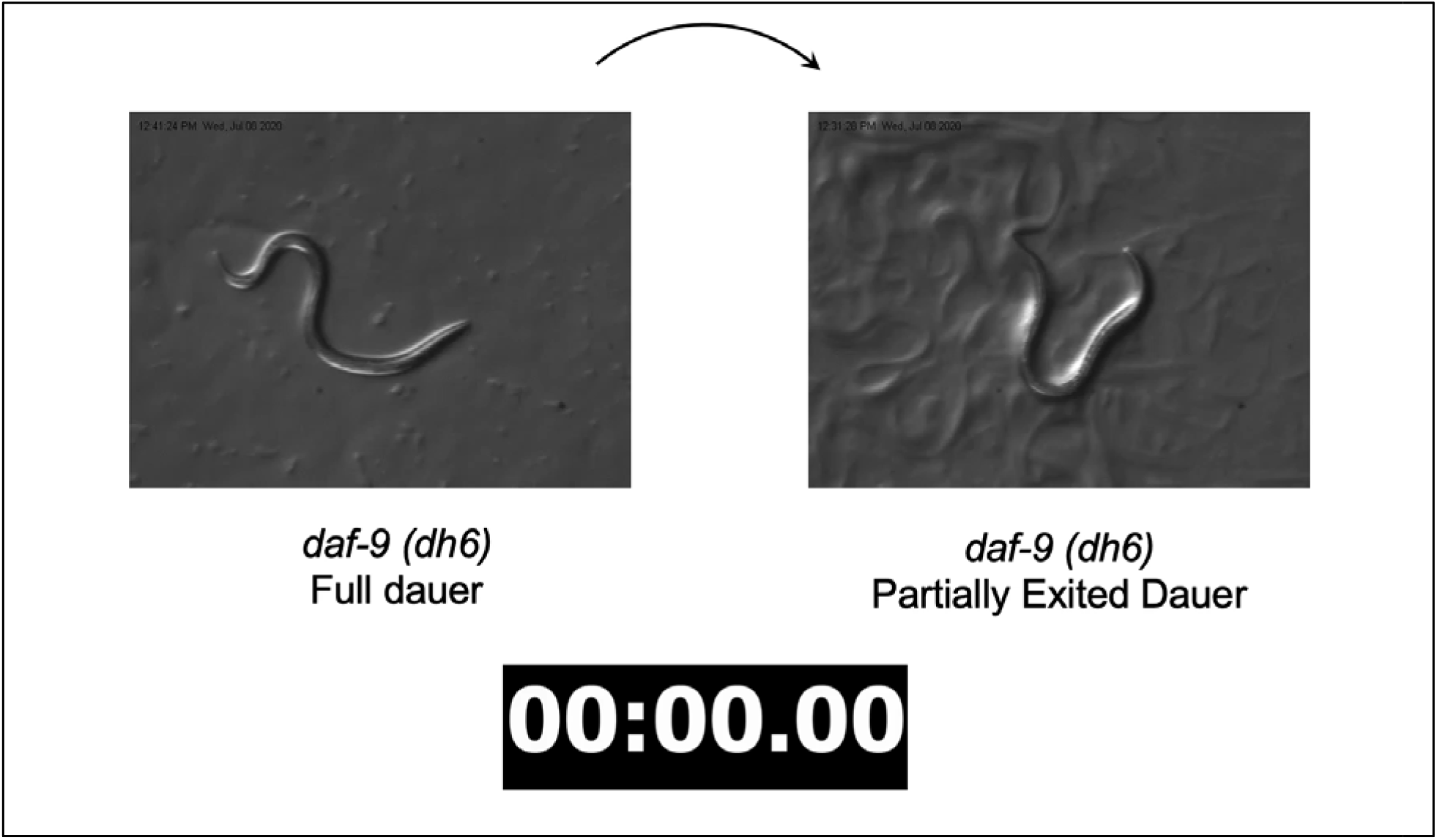
Locomotion behavior of *daf-9(dh6)* partially exited dauers. Shown are ten-second recordings of (left) a *daf-9(dh6)* full dauer formed under unfavorable growth conditions and (right) a *daf-9(dh6)* partially exited dauer formed by transferring full *daf-9(dh6)* dauers from unfavorable to favorable conditions for 24 hours. A second partially exited dauer can be seen crawling at high speed in the background midway through the recording. The *daf-9(dh6)* partially exited dauer performs many small head movements and frequently reverses, unlike *daf-9(dh6)* full dauers which remain idle.

## Supplemental Figures and Tables

**Figure S1.**
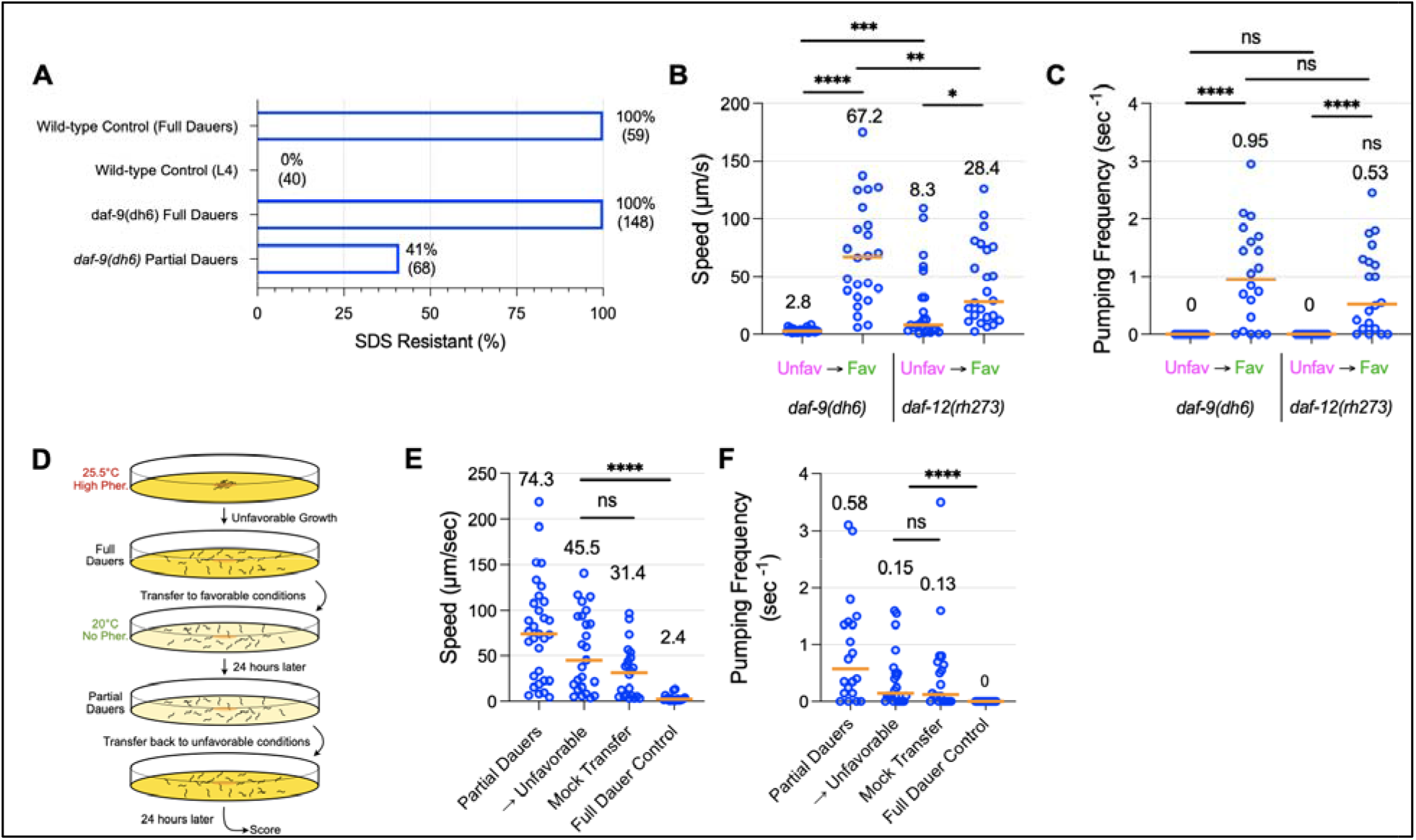
(A) *daf-9(dh6)* full dauers are completely SDS resistant, while *daf-9(dh6)* partial dauers are SDS sensitive. Also shown are wild-type control animals grown under unfavorable conditions to form full dauers or under favorable conditions to form SDS-sensitive L4 larvae. Displayed are the percentages of animals that survived SDS treatment and the corresponding number of animals treated. (B,C) *daf-12(rh273)* phenocopies the *daf-9(dh6)* partial dauer exit phenotype. The Daf-c allele *daf-12(rh273)* shows a similar partial dauer exit phenotype to *daf-9(dh6)* as measured by locomotion speed and pumping frequency, as *daf-12(rh273)* full dauers formed under unfavorable conditions were induced to become partially exited dauers that (B) moved faster and (C) pumped more frequently following transfer to favorable conditions. Note that *daf-12(rh273)* full dauers tend to be slightly more mobile than wild-type or *daf-9(dh6)* dauers, despite being fully pumping quiescent. (D-F) The partial dauer exit phenotype is not reversible. (D) Experimental schematic for reversibility assay. *daf-9(dh6)* partial dauers were produced by first inducing full dauers under unfavorable growth conditions followed by transfer to favorable conditions for 24 hours (Partial Dauers). Partially exited dauers were then transferred back to unfavorable conditions for a further 24 hours (→Unfavorable) and assessed for full dauer characteristics, including locomotion and pumping frequency. A 24-hour incubation under unfavorable conditions did not significantly decrease (E) locomotion speed nor (F) pumping quiescence compared to a mock transfer control, in which partially exited dauers were transferred to favorable conditions for 24 hours. ns, not significant. *, p<0.05. ***, p<0.01. ****, p<0.001 by Mann Whitney test. Each dot is one animal. n ≥ 12 animals for each experimental sample.

**Figure S2:**
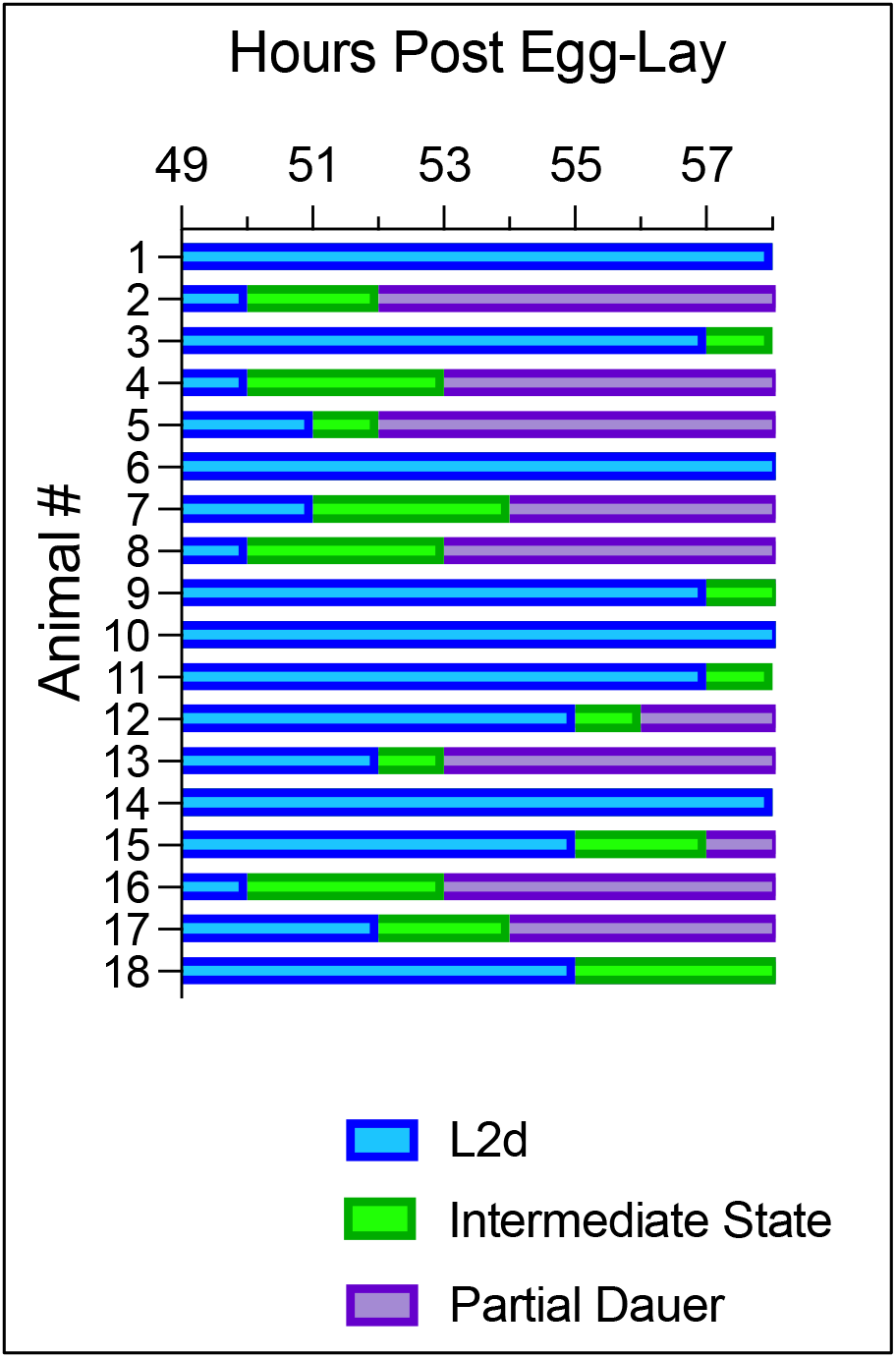
*daf-9(dh6)* mutants were not observed to enter a full dauer state at lower temperatures. *daf-9(dh6)* mutants were grown in absence of exogenously added pheromone at 20.0°C. At 49 hours post egg-lay, animals were individually transferred to new plates and observed every hour. Animals could not be found to pass through a full dauer state as they did at 25.5°C. Instead, following the L2d molt, they passed through an intermediate state that showed characteristics between those of an L2d and a partial dauer larva (see Main Text).

**Figure S3:**
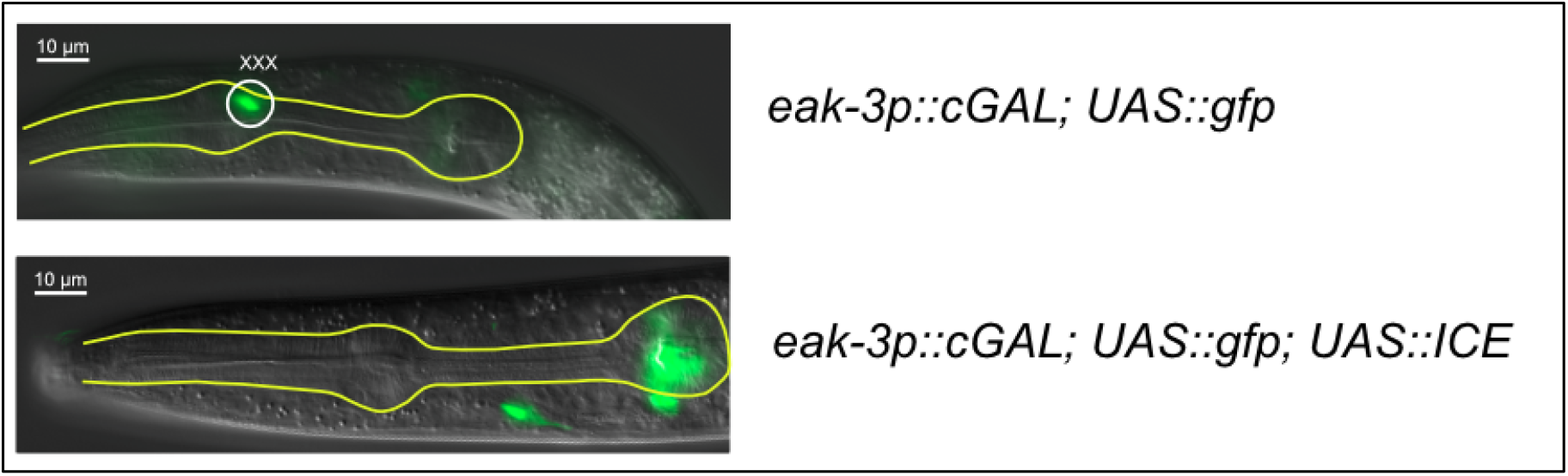
Genetic ablation of the XXX neuroendocrine cells using the human caspase ICE gene. (Top) A transgenic strain using the cGAL bipartite gene expression system expresses *UAS::gfp* using the cell-specific driver *eak-3p::cGAL* in the XXX neuroendocrine cells. The XXX cells are located near the anterior bulb of the pharynx. (Bottom) Expression of the human caspase gene *ICE* using the same XXX-specific *eak-3* cGAL driver shows a loss of fluorescence in the XXX cells. The labeled neuron in the bottom image comes from RFP bleed-through from a co-injection marker labeling the AIY neuron.

**Table S1:**
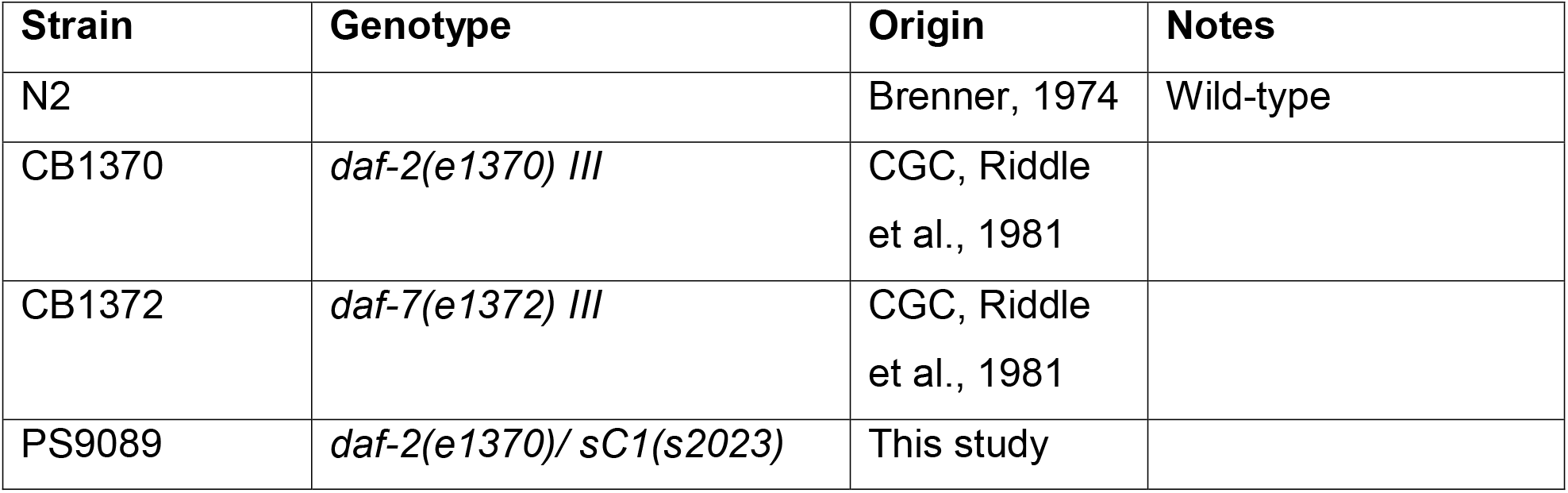
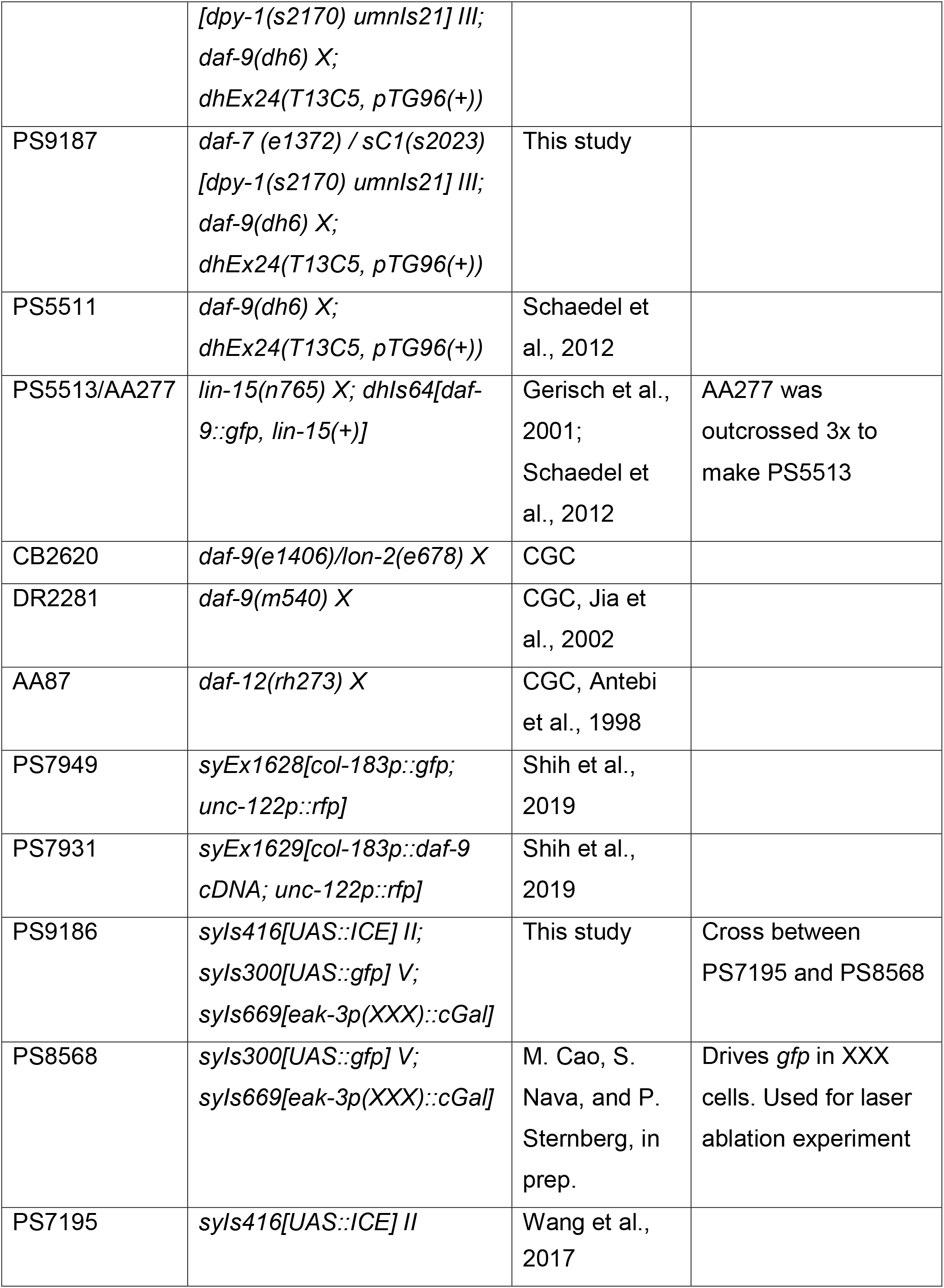
Strain names, genotypes, and origins.

